# Frequency-Dependent Structural Responses of Amyloid- Fibrils to Terahertz Pulses: A Nonequilibrium Molecular Dynamics Study

**DOI:** 10.64898/2026.11.06.764785

**Authors:** Yunyu Wang, Jin Dai, Xubiao Peng, Qing Zhao

**Affiliations:** Center for Quantum Technology Research, Key Laboratory of Advanced Optoelectronic Quantum Architecture and Measurements (MOE), School of Physics, Beijing Institute of Technology, Beijing 100081, China

**Keywords:** Amyloid fibrils, nonequilibrium molecular dynamics, terahertz pulses, mid-infrared pulses, *β*-sheet stability, hydrogen-bond dynamics

## Abstract

The abnormal aggregation of amyloid proteins into fibrils is closely associated with several neurodegenerative diseases, highlighting the importance of developing effective strategies to modulate fibril structural stability. Here, we employed nonequilibrium molecular dynamics(NEMD) simulations to examine the effects of intense terahertz-to-mid-infrared pulses on three amyloid fibrils (2LMP, 2MXU, and 5OQV), focusing on the combined effects of pulse frequency and electric-field strength. Frequency scans from 0.5 to 60 THz in 0.5 THz steps revealed frequency-dependent losses of *β*-sheet content, with several common response patterns in these fibrils. Comparisons at 2, 23, 33, and 50 THz showed that pulse-induced changes in secondary structure differed from those in high-temperature controls. Hydrogen-bond analysis showed that total hydrogen-bond counts could partially recover even as interchain hydrogen bonds continued to decrease, with distinct responses among representative chain pairs. We therefore propose a two-threshold electric-field framework for fibril disruption: *E*_th1_ describes the onset of structural perturbation, whereas *E*_th2_ indicates persistent structural loss after the pulse. This framework offers a testable approach for characterizing amyloid structural responses to external fields, although the underlying energy-coupling mechanisms, quantitative thresholds, and long-term recovery require further validation. These findings may provide a theoretical basis for exploring external-field-based strategies to modulate protein aggregation.

## 1 Introduction

Alzheimer’s disease (AD) is a neurodegenerative disorder of the central nervous system and a major cause of dementia. According to the World Health Organization (WHO), more than 55 million people worldwide were living with dementia, and this number is projected to reach 139 million by 2050 (*1*, *2*). The two main pathological hallmarks of AD are extracellular amyloid plaques formed by the accumulation of amyloid-*β* (A*β*) peptides and intracellular neurofibrillary tangles (NFTs) composed of hyperphosphorylated tau protein (*3*, *4*). A*β* peptides are produced by the sequential cleavage of amyloid precursor protein (APP) by *β*- and *γ*-secretases. The two major forms, A*β*40 and A*β*42, have a strong tendency to aggregate (*5*, *6*). Soluble A*β* oligomers are considered major neurotoxic species that impair synaptic function, while A*β* fibrils can further accumulate into insoluble extracellular plaques (*7*, *8*).

Structural disruption of amyloid fibrils does not necessarily indicate complete disaggregation or reduced toxicity. The disaggregation process may also generate cytotoxic oligomers (*9*). Therefore, when evaluating the effects of external physical stimuli on A*β* fibrils, it is important to distinguish changes in secondary structure, overall conformation, and interactions between peptide chains. A*β* accumulation can activate microglia and promote chronic inflammation, leading to neuronal damage. It can also disrupt calcium homeostasis and mitochondrial function, potentially triggering apoptosis (*10* –*12*). Amyloid fibrils formed both in vivo and in vitro typically exhibit a cross-*β*-sheet structure stabilized by a network of backbone hydrogen bonds (HBs). This structure contributes to their high stability and low solubility, making them difficult to remove and posing challenges for the treatment of neurodegenerative diseases (*13*, *14*). In addition, A*β* fibrils exhibit significant structural polymorphism. Even fibrils formed from the same A*β* peptide sequence can adopt different conformations and packing arrangements (*15*). These structural differences may affect fibril stability and conformational responses to external perturbations. Therefore, comparing A*β* fibrils with different structures may help identify common features of their responses to external fields.

Under physiological conditions, tau protein stabilizes microtubules and supports axonal transport (*16*, *17*). However, an imbalance between kinase and phosphatase activities can lead to tau hyperphosphorylation, reducing its ability to bind to microtubules. Hyperphosphorylated tau can then aggregate into paired helical filaments (PHFs), which form NFTs within neuronal cell bodies (*18*). These aggregates disrupt the neuronal cytoskeleton and impair axonal transport, affecting the transport of nutrients and signaling molecules (*19*). The distribution of NFTs forms the basis of Braak staging and is associated with dementia severity (*20*).

Current treatment strategies for Alzheimer’s disease (AD) still face several challenges. For example, although the anti-amyloid-*β* (A*β*) monoclonal antibody aducanumab has shown therapeutic potential, the clinical significance of its reported benefits remains debated (*21*). Small-molecule drugs also face difficulties in crossing the blood–brain barrier (*22*). Physical approaches have limitations as well. Focused ultrasound has been investigated to promote microglia-mediated clearance of A*β*, but concerns remain regarding potential tissue damage and limited molecular selectivity (*23*). Gene-based therapies also face challenges in achieving efficient delivery to the central nervous system (*24*). Overall, existing strategies still face challenges in selectivity, delivery, and safety (*25*).

THz radiation lies between the microwave and infrared regions of the electromagnetic spectrum, typically spanning 0.1–10 THz. Its photon energy is relatively low, approximately 4.1 meV at 1 THz (*26*, *27*). THz fields can interact with protein collective motions and intermolecular interactions, offering a potential approach to modulating protein structure without chemical reagents (*28*, *29*) and the solvent environment may also influence protein responses to external fields. Sugiyama et al. showed that sub-THz irradiation could alter coupled protein–water dynamics and protein hydration (*30*) which highlighted the importance of comparing structural responses to external fields at different frequencies with those caused by conventional heating, while examining changes in intermolecular interactions.

Early studies by Govorun et al. suggested that far-infrared radiation could affect protein structure and stability (*31*). Homenko et al. reported that irradiation at 0.1 THz and 0.08 mW*/*cm^2^ slightly reduced alkaline phosphatase activity and decreased the stability of preformed antigen–antibody complexes (*32*). More recently, Zhang et al. demonstrated that 0.14 THz irradiation reduced A*β* deposition and tau hyperphosphorylation in AD mice, although the underlying molecular mechanisms remain unclear (*33*). THz spectroscopy has also revealed long-range protein vibrations, providing evidence for the interaction of low-frequency electromagnetic fields with collective protein motions (*34*).

Research has extended beyond the low-frequency THz region to higher frequencies, including the 20–60 THz mid-infrared range (*35*, *36*). In this region, the protein amide I band typically lies at 1600–1700 cm*^−^*^1^, corresponding to approximately 48–51 THz (*37*). Kawasaki et al. reviewed infrared free-electron laser (IR-FEL) experiments demonstrating the disassembly of several amyloid fibrils, including lysozyme fibrils, although the detailed mechanisms require further investigation (*38*). Using NEMD simulations, Hoang Viet et al. showed that resonant excitation of amide I vibrations by infrared laser pulses could disrupt amyloid fibrils (*39*). Related experimental and simulation studies have further examined the relationship between amide I excitation and fibril dissociation (*40*). In addition, Peng et al. reported that irradiation at 34.88 THz slowed A*β* oligomer fibrillization, reducing the aggregation rate to approximately 80% of the untreated control (*35*).

Based on these findings, we performed NEMD simulations over 0.5–60 THz at 0.5 THz intervals to investigate the structural responses of A*β* fibrils to low-frequency THz and midinfrared pulses. Three A*β* fibril models with distinct packing arrangements were examined under intense oscillating electric fields. Frequency-dependent structural changes were characterized by analyzing *β*-sheet content, secondary-structure composition, radius of gyration (*R_g_*), solvent-accessible surface area (SASA), and interchain hydrogen-bond dynamics. At selected frequencies, we compared pulse-induced structural changes with those under high-temperature conditions and examined local conformational changes and interchain interactions in different regions of the fibrils. We further varied the peak electric-field strength to investigate the onset of structural perturbation and persistent *β*-sheet loss after the pulse, proposing a two-threshold framework (*E*_th1_/*E*_th2_) to characterize these responses. Together, these analyses provide molecular-level insights into the structural responses of A*β* fibrils to intense oscillating electric fields.

## 2 Results

To investigate the effects of intense pulsed electric fields on A*β* fibril stability, we first examined the structural responses of three fibril models to a 50 THz mid-infrared pulse using NEMD simulations. Figure 1(a) shows the initial and post-pulse structures of the three models. The applied pulse had a peak electric-field strength of 2 V*/*nm, with fieldfree simulations (*E*_0_ = 0) serving as controls. As illustrated in Figure 1(b), the electric field was polarized along the *x*-axis, with the *z*-axis representing the nominal propagation direction. A spatially uniform, time-dependent electric field was applied without explicitly modeling electromagnetic wave propagation.

**Figure 1:**
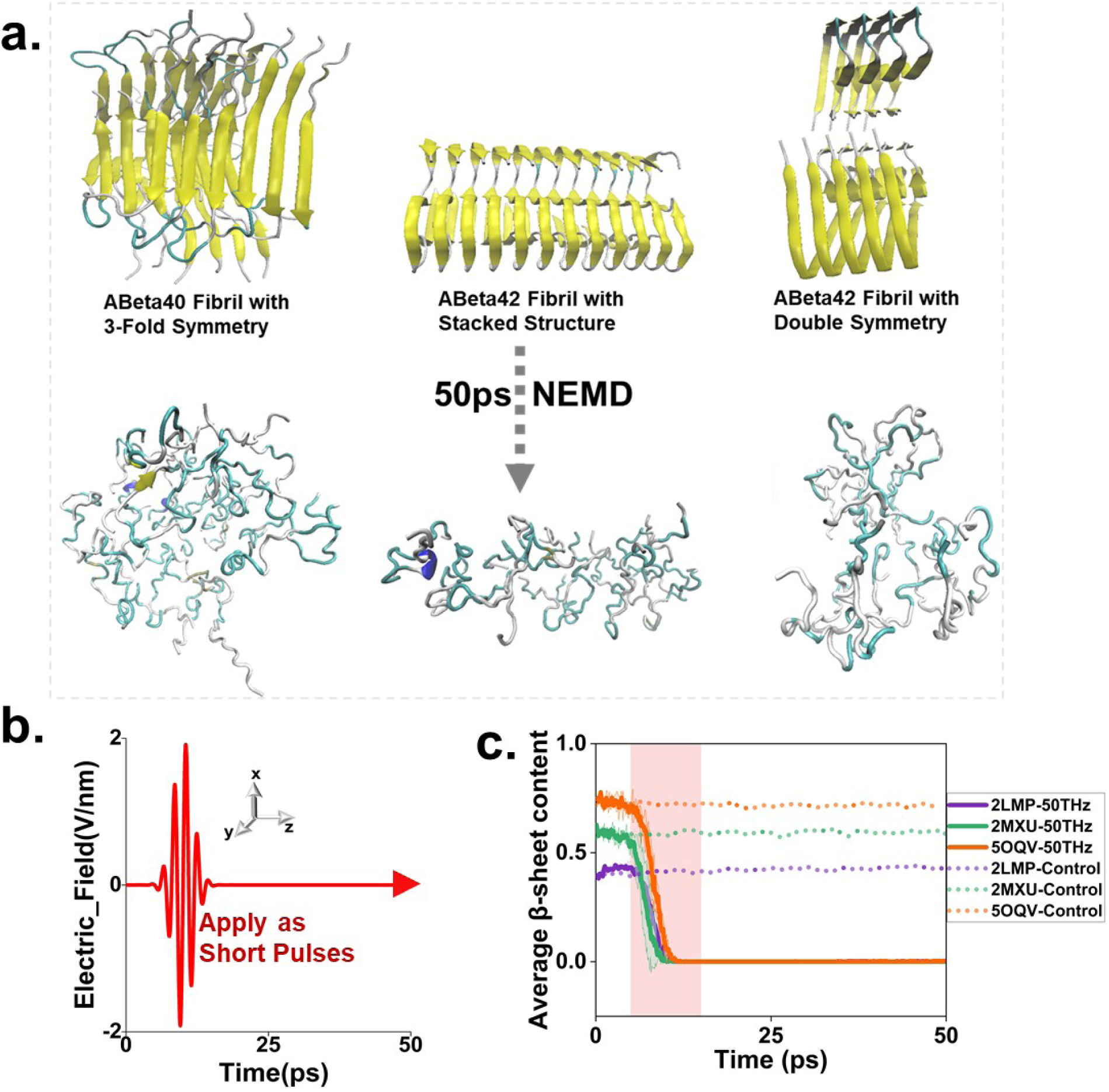
Pulse-induced structural changes in three A*β* fibril models. (a) Initial structures (top) and structures after exposure to a 50 THz pulse (bottom) for 2LMP, 2MXU, and 5OQV. (b) Schematic of the Gaussian-envelope pulse at 50 THz with a peak electric-field strength of 2 V*/*nm. The electric field is polarized along the *x*-axis, with the *z*-axis indicating the nominal propagation direction. (c) Average *β*-sheet content of the three fibril models under the 50 THz pulse (solid lines) and the field-free control (*E*_0_ = 0, dotted lines). Shaded regions around the curves represent the standard deviation from three independent simulations, while the pink background indicates the main pulse excitation period. All three models exhibit substantial *β*-sheet loss, suggesting a common structural response despite differences in fibril architecture.

Figure 1(c) shows the evolution of the average *β*-sheet content over 50 ps. All three fibril models exhibited substantial reductions in *β*-sheet content under the 50 THz pulse compared with their field-free controls. These changes occurred mainly during the pulse excitation period, indicated by the pink shaded region, although the Gaussian envelope remained nonzero outside this interval. The similar responses across the three models suggest that *β*-sheet destabilization can occur in fibrils with different packing arrangements. These findings motivated a systematic frequency scan from 0.5 to 60 THz to further examine the frequency dependence of fibril structural stability (Figure 2).

**Figure 2:**
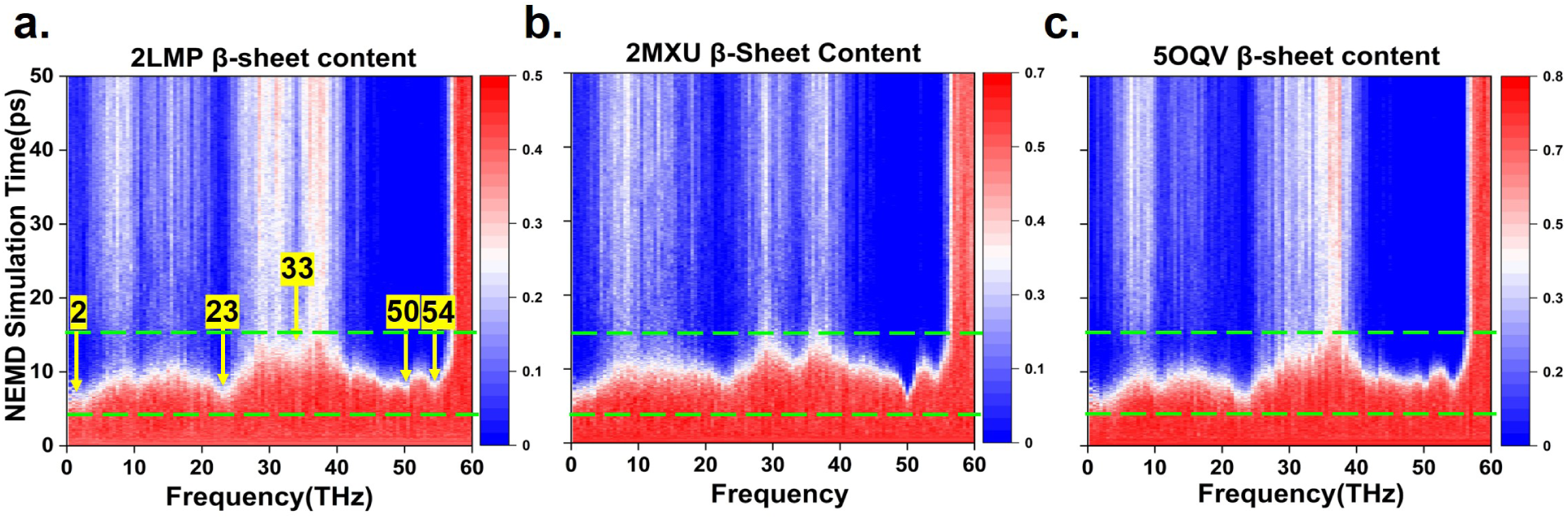
Frequency-dependent structural responses of three A*β* fibril models. (a–c) Average *β*-sheet content of 2LMP, 2MXU, and 5OQV, respectively, over 120 nonzero frequencies from 0.5 to 60 THz at 0.5 THz intervals. The peak electric-field strength was 2 V*/*nm, with polarization along the *x*-axis and a simulation duration of 50 ps. The horizontal axis represents frequency (THz). Red and blue indicate higher and lower *β*-sheet content, respectively, with different upper limits for the color scales of the three models. Green dashed lines mark the main pulse excitation period (approximately 5–15 ps), although the Gaussian envelope remains nonzero outside this interval. Yellow arrows indicate 2, 23, 33, 50, and 54 THz.

### 2.1 Frequency-Dependent Effects of THz and Mid-Infrared Pulses on Amyloid Fibril *β*-Sheet Stability

Following the substantial *β*-sheet loss observed in all three A*β* fibril models under the 50 THz pulse (Figure 1), we performed a systematic frequency scan from 0.5 to 60 THz at 0.5 THz intervals to examine the frequency dependence of fibril structural stability.

Figure 2a–c shows the results for 2LMP, 2MXU, and 5OQV, respectively. All simulations were performed with a peak electricfield strength of 2 V*/*nm over 0–50 ps. The heatmaps show the average *β*-sheet content as a function of frequency and time, with red and blue indicating higher and lower contents, respectively. Green dashed lines mark the main pulse excitation period. Different color scales were used to account for differences in the initial *β*-sheet content among the models.

Despite their different packing arrangements, all three fibril models exhibited frequency-dependent *β*-sheet loss across the scanned range. Although the magnitude of the response varied with frequency, several similar response patterns were observed among the models.

To assess whether *β*-sheet loss was accompanied by changes in overall fibril conformation, we calculated the radius of gyration (*R_g_*) across the frequency range (Figure S1, Supporting Information). All three models showed increased *R_g_* at selected frequencies, particularly near 50 THz. These changes were consistent with the *β*-sheet loss in Figure 2, indicating that secondary-structure disruption was accompanied by changes in overall fibril conformation.

Based on the frequency scan, we selected 2, 23, 33, and 50 THz for further analysis. The 2 THz frequency represents the low-frequency THz region, while 23 and 33 THz lie outside the amide I region. The 50 THz frequency corresponds to approximately 1668 cm*^−^*^1^, within the typical protein amide I band (*37*). At the applied field strength, pronounced *β*-sheet loss was observed at 2, 23, and 50 THz, whereas more ordered structure was retained at 33 THz. Strong structural responses were also observed near 50 and 54 THz, while *β*-sheet loss was relatively limited near 60 THz. These results indicate a nonmonotonic dependence of structural disruption on pulse frequency. Although similar structural changes were observed at 50 and 54 THz (Figure S2, Supporting Information), this similarity alone does not establish that the two frequencies excite the same vibrational mode.

Given the shared response features among the three models, 2LMP was selected for further analysis. Figure 3 compares the time evolution of secondary-structure components under pulses at 2, 23, 33, and 50 THz, together with field-free controls at 310 and 500 K. At 2, 23, and 50 THz, the *β*-sheet content decreased substantially, accompanied by a rapid increase in irregular structures, some of which subsequently transitioned into bends or turns. In contrast, more *β*-sheet structure was retained at 33 THz. The 310 K control remained relatively stable over 100 ns, whereas the 500 K control showed progressive *β*-sheet loss over 200 ns.

**Figure 3:**
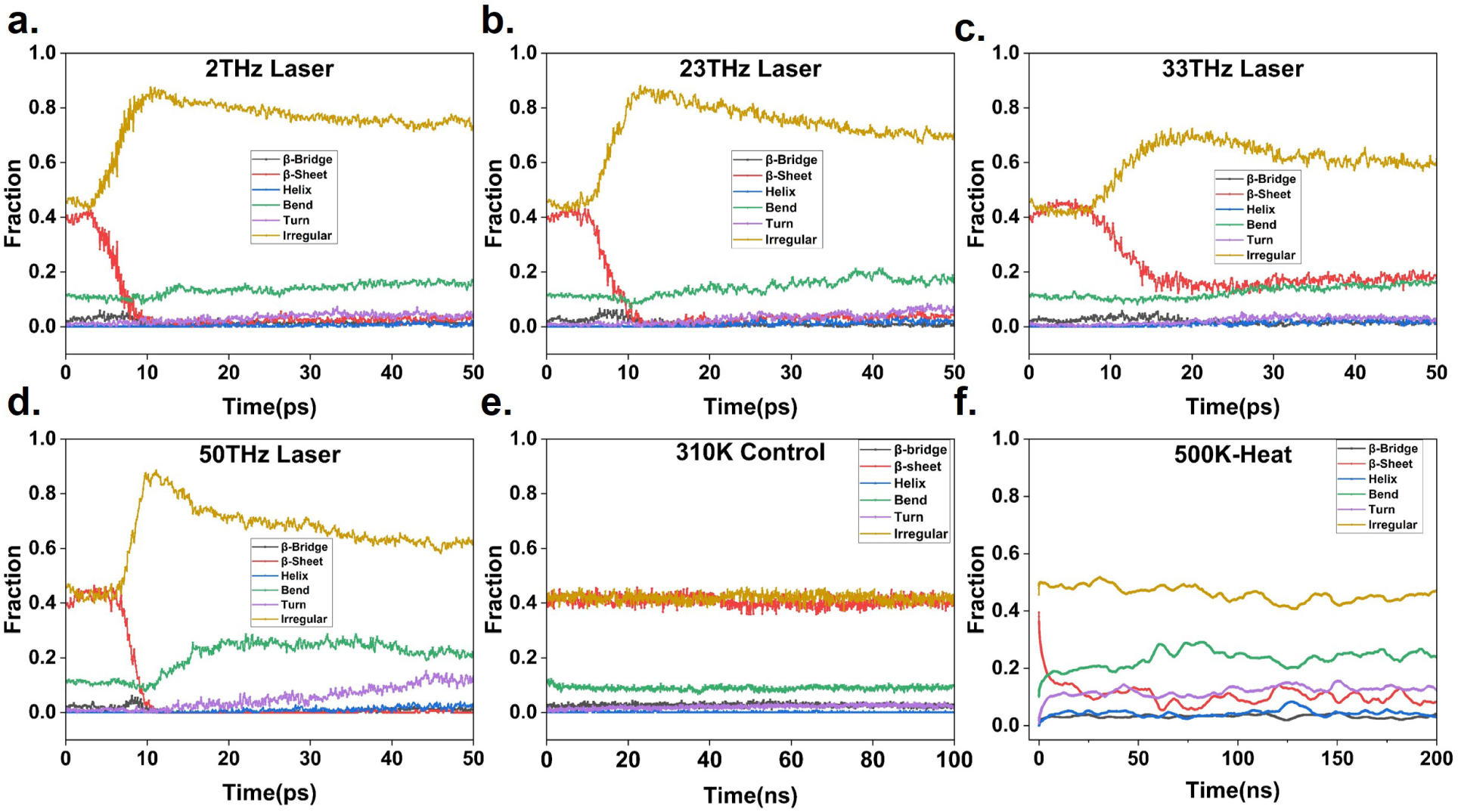
Time evolution of secondary-structure components in the 2LMP fibril under pulsed-field excitation and temperature controls. (a–d) Pulses at 2, 23, 33, and 50 THz, respectively, with a peak electric-field strength of 2 V*/*nm, *x*-polarization, and a simulation duration of 50 ps. (e) Field-free control at 310 K for 100 ns. (f) Field-free high-temperature control at 500 K for 200 ns. Gray, red, blue, green, purple, and gold curves represent *β*-bridges, *β*-sheets, helices, bends, turns, and irregular structures, respectively. Panels are arranged from left to right and top to bottom. Time is shown in ps for (a–d) and ns for (e,f). All panels show time-dependent secondary-structure content rather than endpoint distributions. The high-temperature control was not matched to the absorbed pulse energy.

The redistribution of secondary-structure components differed between pulse excitation and high-temperature conditions. Under the 500 K control, the loss of *β*-sheet structure was mainly accompanied by increases in bends and turns, with only a slight increase in irregular structures. In contrast, pulse excitation produced a more pronounced increase in irregular structures, despite less extensive *β*-sheet loss under the examined conditions. These observations suggest different pathways of structural evolution.

To further examine the relationship between secondary-structure changes and solvent exposure, we analyzed the total solvent-accessible surface area (SASA) of 2LMP at selected frequencies (Figure S3, Supporting Information). Total SASA increased to varying degrees at all selected frequencies, with particularly pronounced increases at 50 and 54 THz. SASA continued to rise after the main pulse excitation period and then partially decreased, remaining above its initial value at 50 ps. These results indicate that pulse-induced *β*-sheet loss was accompanied by increased solvent exposure and conformational rearrangement.

### 2.2 THz Pulse-Induced Targeted Disruption of Hydrogen-Bond Networks in Amyloid Fibrils

The preceding results show that intense pulses at different frequencies induced varying degrees of *β*-sheet loss and secondary-structure rearrangement in A*β* fibrils (Figure 3). Given the key role of hydrogen bonds in stabilizing ordered *β*-sheet structures, we further analyzed hydrogen-bond dynamics in 2LMP under different pulse conditions and compared them with the 500 K high-temperature control (Figure 4).

**Figure 4:**
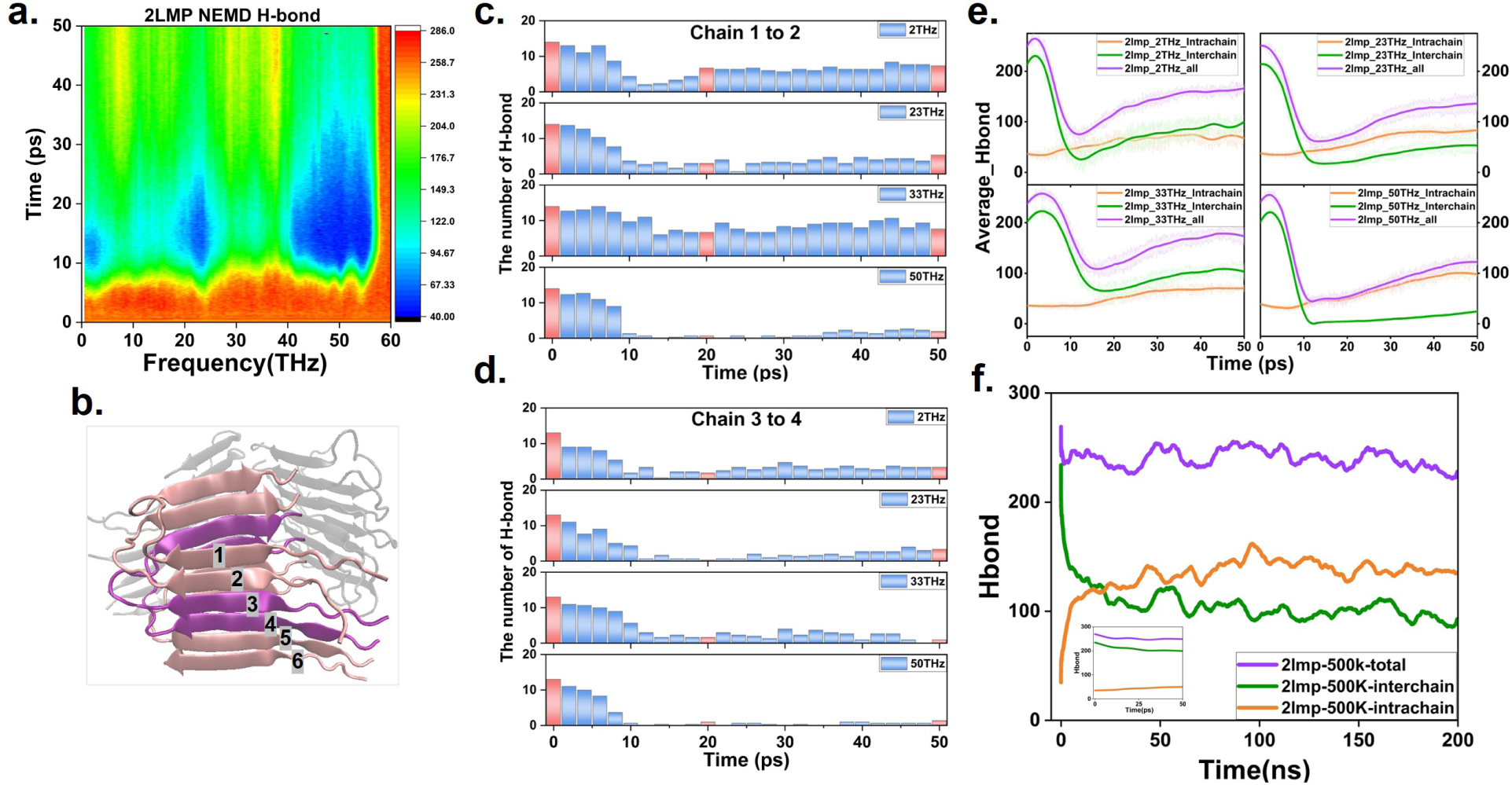
Hydrogen-bond network responses of the 2LMP fibril. (a) Heatmap of the total hydrogen-bond count across 0.5–60 THz, with a peak electric-field strength of 2 V*/*nm, polarization along the *x*-axis, and a simulation duration of 50 ps. (b) Structural locations of representative chain pairs. Chains 1–2 are located near the outer region of the fibril stack, whereas chains 3–4 lie closer to the interior. (c,d) Time evolution of hydrogen-bond counts for chain pairs 1–2 and 3–4, respectively. The panels in each column correspond to 2, 23, 33, and 50 THz from top to bottom. (e) Time evolution of intrachain (orange), interchain (green), and total (purple) hydrogen-bond counts at the four representative frequencies. (f) Corresponding hydrogen-bond counts under the 500 K high-temperature control over 200 ns.

Figure 4a shows the time evolution of the total hydrogen-bond count in the 2LMP fibril across the 0.5–60 THz frequency range. Pulse-induced hydrogen-bond loss exhibited a clear frequency dependence. At 50 THz, the total hydrogen-bond count decreased rapidly during the main excitation period, followed by partial recovery. The extent of hydrogen-bond loss and subsequent recovery varied at other frequencies. These results indicate that the applied field frequency affects both hydrogen-bond disruption and subsequent network reorganization.

At 2, 23, and 33 THz, hydrogen-bond disruption and recovery varied in magnitude. Figure 4b illustrates the spatial locations of representative chain pairs: chains 1–2 are located near the outer region of the fibril stack, whereas chains 3–4 lie closer to the interior. Figures 4c and 4d show that intense THz pulses preferentially disrupt interchain hydrogen bonds in the fibril interior, with a significantly stronger effect on inner than outer chain pairs. At 33 THz, the inner chain pair retained substantially fewer hydrogen bonds than the outer pair, while the 50 THz pulse caused extensive disruption in both pairs. These results demonstrate spatially heterogeneous responses, with preferential disruption of the fibril interior, in contrast to the behavior under thermal conditions.

To extend the analysis, we examined interchain hydrogen-bond dynamics across three groups of chain pairs (Figure S4, Supporting Information). At 50 THz, interchain hydrogen bonds decreased rapidly in all groups during pulse excitation, followed by limited recovery. At other frequencies, the extent of hydrogen-bond reorganization varied among the groups.

Figure 4e quantifies the changes in intrachain, interchain, and total hydrogen-bond counts in 2LMP at the four representative frequencies. At effective frequencies such as 50 THz, the total hydrogen-bond count decreased rapidly during pulse excitation, followed by partial recovery. This recovery mainly resulted from the reformation of intrachain hydrogen bonds, whereas interchain hydrogen bonds remained at low levels after the pulse. These results indicate that the interchain interaction network was severely disrupted and could not recover spontaneously. Thus, the partial recovery of total hydrogen bonds does not indicate restoration of the original interchain network. The reformation of intrachain hydrogen bonds marks irreversible separation between peptide chains, indicating the irreversible nature of THz pulse-induced fibril disaggregation. Together with the persistent *β*-sheet loss shown in Figure 3, these findings demonstrate that disruption of the ordered fibril structure is accompanied by substantial reorganization of the interchain interaction network.

To compare hydrogen-bond dynamics under pulse excitation and high-temperature conditions, Figure 4f shows the changes in intrachain, interchain, and total hydrogen-bond counts in 2LMP at 500 K. During the initial 0–50 ps, hydrogen-bond changes in the high-temperature control were relatively gradual, without the rapid interchain hydrogen-bond loss observed under intense pulse excitation. With prolonged heating, interchain hydrogen bonds gradually decreased, while intrachain hydrogen bonds increased and continued to fluctuate on the nanosecond timescale. Figure S5 (Supporting Information) further shows the hydrogen-bond dynamics of three representative chain-pair groups under high-temperature conditions, revealing differences in hydrogen-bond retention and reorganization among interchain interfaces. These results show that both intense pulses and high-temperature heating induce hydrogen-bond network reorganization, but with distinct structural evolution and timescales.

Overall, the hydrogen-bond analysis shows that pulse-induced *β*-sheet loss is accompanied by rapid disruption of the interchain hydrogen-bond network and subsequent intrachain hydrogen-bond reorganization. Differences among chain-pair responses further suggest that local fibril packing environments may influence hydrogen-bond stability. These findings provide a molecular basis for understanding field-induced destabilization of ordered fibril structures and the associated rearrangement of intermolecular interactions.

### 2.3 Local Geometric Changes and Solvent Exposure in Amyloid Fibrils under THz Pulses

To examine spatial differences in local conformational responses, we analyzed per-residue bond-angle changes and SASA in chain 3 of the 2LMP fibril under a 50 THz pulse (Figure 5). In Figure 5a, bond-angle changes at several residues near the purple vertical dashed line began at similar times, indicating approximately synchronous response onset within the available temporal resolution. However, the duration of pronounced fluctuations varied among residues and generally increased with residue index, indicating longer relaxation times after pulse excitation and distinct local conformational adjustments.

**Figure 5:**
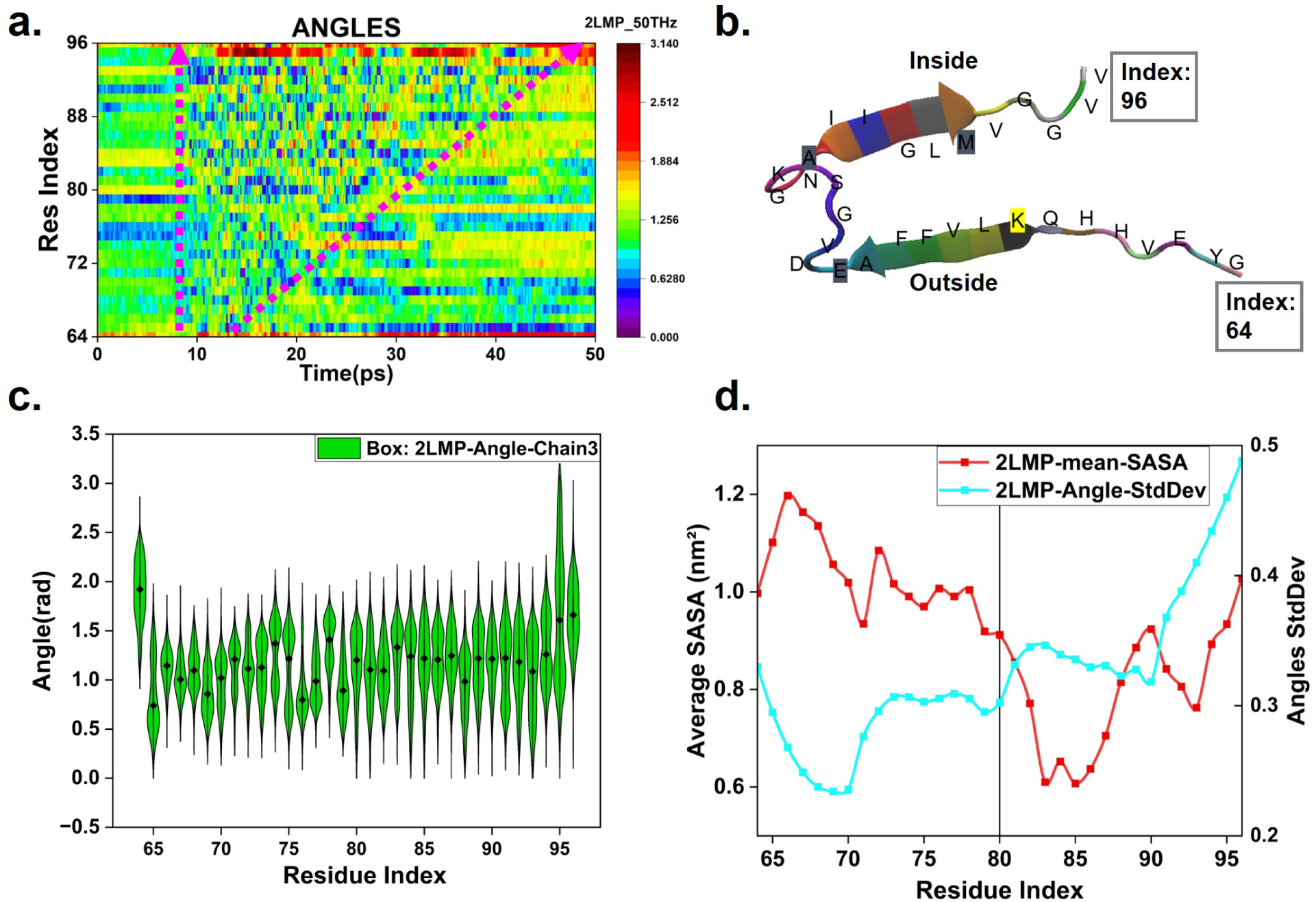
Preliminary analysis of local geometry and solvent exposure in chain 3 of the 2LMP fibril under a 50 THz pulse with a peak electric-field strength of 2 V*/*nm and *x*-polarization. (a) Time-resolved heatmap of Bond-angle, with angles expressed in radians. (b) Conformation of chain 3 with residue indices. Inside and outside indicate the approximate structural locations. (c) Bond-angle distributions at individual positions over 0–50 ps. (d) Mean per-residue SASA (red, left axis, nm^2^) and bond-angle standard deviation (cyan, right axis, rad).

The region marked by the two purple dashed arrows in Figure 5a shows frequent color alternations, reflecting substantial bond-angle fluctuations over time. These changes persisted longer at higher residue indices. As illustrated in Figure 5b, these residues are mainly located in chain segments facing the fibril interior. This indicates that inner and outer chain segments exhibit different local geometric responses to the pulse in both magnitude and duration.

Figure 5c further presents violin plots of the bond-angle distributions for individual residues over the analysis period. A broader angular distribution with longer tails indicates a larger range of bond-angle variations, while the width of each violin represents the distribution density at a given angle. The distributions varied among residues, with some higher-index residues showing broader distributions, consistent with the persistent fluctuations observed in Figure 5a. Figure 5d further quantifies these variations using the bond-angle standard deviation and compares them with the mean per-residue SASA.

Some inward-facing chain segments exhibited relatively low SASA but large bond-angle fluctuations, whereas certain outward-facing segments showed greater solvent exposure with smaller angular variations. These results indicate that the magnitude of local conformational changes does not depend solely on solvent exposure but may also be influenced by the local packing environment and intermolecular constraints. To further examine this behavior, we extended the per-residue SASA and bond-angle analyses to all 18 peptide chains of 2LMP (Figures S6 and S7, Supporting Information). Differences in solvent exposure and geometric fluctuations among chains and residues further demonstrate the spatial heterogeneity of local structural responses.

Combined with the hydrogen-bond changes shown in Figure 4, these findings suggest that pulse-induced reorganization of interchain interactions may alter local structural constraints. Rearrangement of contacts and hydrogen-bond networks in the fibril interior may lead to larger and more persistent bond-angle changes, affecting the magnitude and duration of local conformational adjustments. Thus, local fibril responses may be jointly influenced by solvent exposure, molecular packing, and interchain interactions, providing further insight into pulse-induced structural destabilization.

### 2.4 Effects of THz Pulse Intensity on Amyloid Fibril Disaggregation

To examine the effects of peak electric-field strength on fibril stability, we performed systematic field-strength scans up to 2 V*/*nm at four representative frequencies (2, 23, 33, and 50 THz). Changes in the *β*-sheet content of 2LMP were analyzed over 0–50 ps. Figure 6 presents the field-strength-dependent response heatmaps at each frequency. As the field strength increased, the fibril gradually developed local structural perturbations, followed by substantial and persistent *β*-sheet loss at higher field strengths. These results indicate that the applied field strength influences both the magnitude of structural changes and whether the initial perturbations develop into sustained structural destabilization.

**Figure 6:**
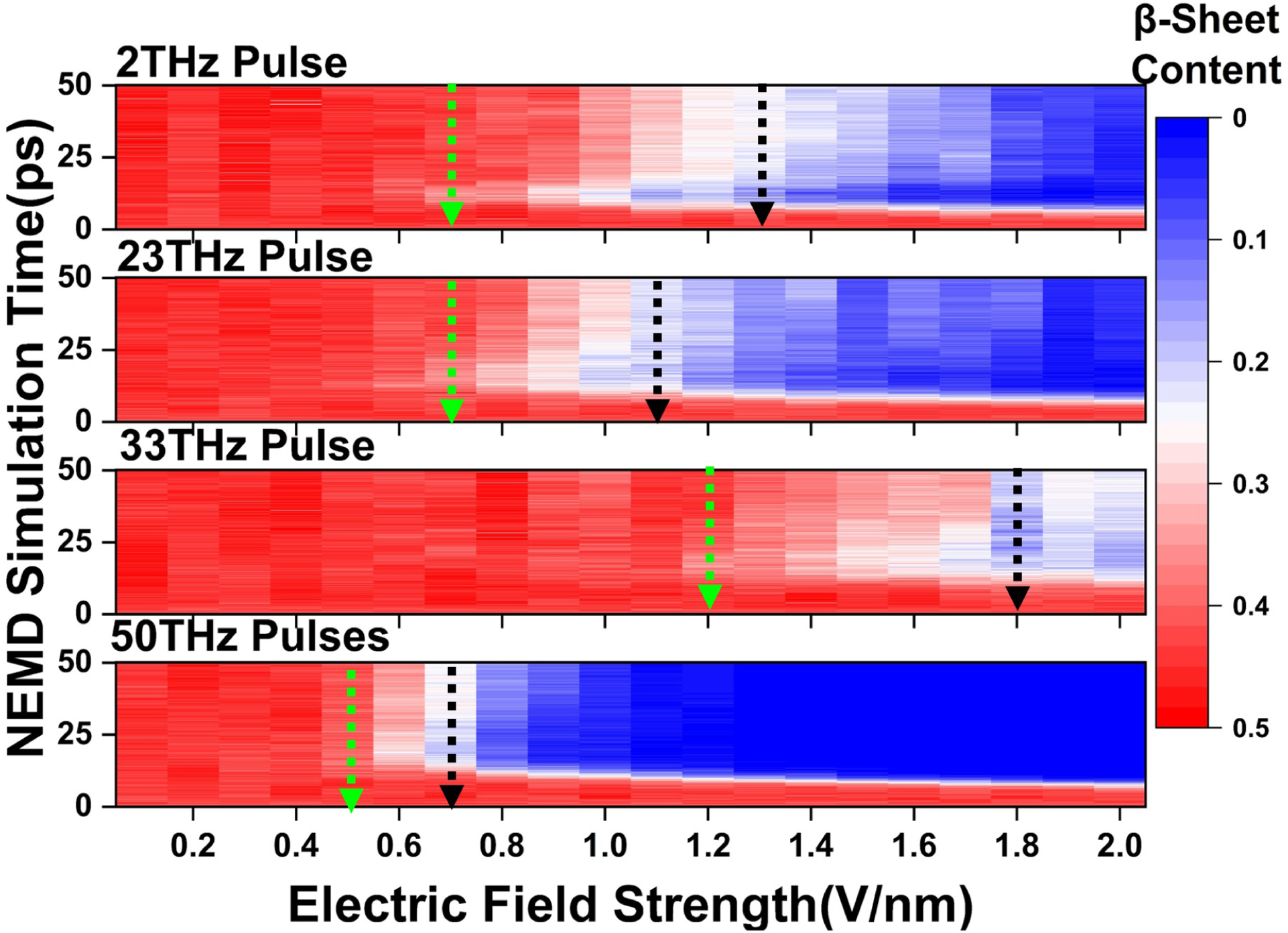
Field-strength-dependent structural responses of the 2LMP fibril at four representative frequencies. From top to bottom, the four rows correspond to 2, 23, 33, and 50 THz, respectively. The horizontal axis represents the peak electric-field strength *E*_0_ (V*/*nm), and the vertical axis represents NEMD simulation time (0–50 ps). The electric field is polarized along the *x*-axis. Red and blue indicate higher and lower *β*-sheet content, respectively. The field-strength range extends to 2 V*/*nm. Green and black arrows indicate the estimated *E*_th1_ and *E*_th2_ thresholds, respectively.

Based on the magnitude and persistence of *β*-sheet loss, we further propose a two-threshold electric-field framework that divides the structural response into three regimes: weak response, transitional response, and sustained destabilization. The first threshold, *E*_th1_ (green arrows in Figure 6), marks the onset of detectable structural perturbation, whereas the second threshold, *E*_th2_ (black arrows), marks the field strength at which substantial *β*-sheet loss begins to persist until the end of the simulation window. Between these thresholds lies the transitional regime, where the extent of structural loss and subsequent recovery varies with pulse frequency and field strength.

As shown in Figure 6, the estimated *E*_th1_ values at 2, 23, 33, and 50 THz were approximately 0.7, 0.7, 1.2, and 0.5 V*/*nm, respectively, while the corresponding *E*_th2_ values were approximately 1.3, 1.1, 1.8, and 0.7 V*/*nm. Among these frequencies, 50 THz exhibited the lowest thresholds for structural perturbation and sustained destabilization, as well as the narrowest transitional regime. At field strengths of approximately 1 V*/*nm or higher, the post-pulse *β*-sheet content approached zero. In contrast, 33 THz showed higher thresholds, with some *β*-sheet structure remaining even at 2 V*/*nm. Although 2 and 23 THz had similar onset thresholds, their thresholds for sustained destabilization differed, indicating distinct field-strength dependencies of structural perturbation and persistent *β*-sheet loss. These results are consistent with the frequency-dependent responses shown in Figures 2 and 3.

Combined with the interchain hydrogen-bond analysis in Figure 4 and Figure S4 (Supporting Information), the observed *β*-sheet loss is consistent with disruption of the hydrogen-bond network that stabilizes the ordered fibril structure. This suggests that field-strength-dependent structural destabilization may involve the reorganization of interchain interactions. The proposed two-threshold framework links pulse frequency to the extent of structural responses, providing a common basis for comparing fibril sensitivity and the field strengths required for structural disruption.

These thresholds are empirical parameters determined under specific pulse conditions and within a 50 ps observation window. They depend on pulse frequency, envelope shape, initial configuration, and observation time. Because electric-field strength and energy have different physical dimensions, neither *E*_th1_ nor *E*_th2_ can be directly interpreted as a hydrogen-bond energy barrier or interchain binding energy. No second threshold is assigned when sustained destabilization is not clearly observed within the scanned field-strength range. Further analysis of individual replicate trajectories is needed to estimate the thresholds and their uncertainties before examining their molecular origins.

## 3 Discussion

Through systematic NEMD simulations, we demonstrate frequency- and field-strength-dependent structural destabilization across three A*β* fibril models with distinct packing arrangements. The results reveal substantial *β*-sheet loss, reorganization of interchain hydrogen-bond networks, and spatially heterogeneous conformational responses. Furthermore, a two-threshold electric-field framework is proposed to characterize the onset of structural perturbation and sustained *β*-sheet loss, providing a systematic basis for evaluating fibril responses to intense oscillating electric fields.

First, despite differences in packing arrangements and peptide lengths, the three A*β* fibril models (2LMP, 2MXU, and 5OQV) exhibited similar *β*-sheet loss at several frequencies, suggesting common factors governing their structural responses. The frequency of 50 THz corresponds to approximately 1668 cm*^−^*^1^, within the typical protein amide I band, allowing comparison with previous studies of mid-infrared laser-induced amyloid fibril dissociation (*37*, *40*). In contrast, 54 THz corresponds to approximately 1801 cm*^−^*^1^, above the typical amide I region. Although both frequencies produced similar structural changes at the applied field strength, this does not establish that they excite the same vibrational modes or share the same energy-transfer pathways. Further mode assignment requires absorption spectra calculated with the employed force field and analysis of local vibrations. At 33 THz, more *β*-sheet structure was retained, indicating a weaker structural response. This suggests that 33 THz may lie in a relatively inactive region of the *β*-sheet backbone vibrational spectrum, where key vibrational modes are less effectively excited.

These findings deepen our understanding of the effects of THz radiation on amyloid fibrils and highlight its potential applications. The proposed combination of 50 THz and a peak electric-field strength above 1 V*/*nm may serve as a potential “optical tweezer”-like approach for selectively disaggregating amyloid fibrils while avoiding thermal damage to surrounding healthy tissues. This approach could offer a new route toward noninvasive treatment strategies for neurodegenerative diseases.

Nevertheless, several limitations remain. First, this study is primarily based on theoretical simulations. Further investigations using two-dimensional THz spectroscopy or normal mode analysis are needed to confirm the coupling between the applied fields and specific vibrational modes. Clinical translation will also require evaluation of the disaggregation efficiency and biological safety of THz pulses in animal models, such as AD mice. The present results do not establish an in vivo therapeutic window or the safety of surrounding tissues. Moreover, the size, reaggregation behavior, and toxicity of aggregates formed after fibril destabilization require independent evaluation. These findings provide a basis for selecting pulse frequencies and field strengths in future experimental studies.

In summary, this study integrates frequency-dependent structural responses, hydrogen-bond network dynamics, and electric-field-strength effects to demonstrate the potential of external-field control over amyloid fibril stability. The proposed frequency–field-strength control framework and inside-out disaggregation mechanism provide new insights into the biophysical effects of THz radiation and offer a theoretical basis for exploring novel therapeutic strategies against neurodegenerative diseases.

## 4 Conclusion

In this study, we systematically investigated the effects of intense THz-to-mid-infrared pulses (0.5–60 THz) on three A*β* fibril models with distinct packing arrangements using NEMD simulations. The structural responses showed a strong dependence on pulse frequency and electric-field strength. Although the three models exhibited similar *β*-sheet loss at selected frequencies, the extent of structural disruption varied, suggesting common response features influenced by molecular packing.

Further structural analyses revealed that pulse-induced *β*-sheet loss was accompanied by increased irregular structures, changes in overall fibril conformation, and substantial disruption of interchain hydrogen-bond networks. After pulse excitation, intrachain hydrogen bonds partially reformed, whereas the original interchain network and ordered *β*-sheet structures did not fully recover within the observation window. Comparisons with high-temperature controls also revealed distinct patterns of secondary-structure evolution and hydrogen-bond reorganization under different perturbation conditions.

Based on these findings, we proposed a two-threshold electric-field framework to distinguish the onset of structural perturbation from sustained destabilization. Among the representative frequencies examined, 50 THz exhibited the lowest estimated thresholds, highlighting the importance of frequency selection in modulating fibril stability.

Overall, this study establishes a relationship among pulse frequency, electric-field strength, and the structural responses of A*β* fibrils at multiple molecular levels. These findings provide insights into field-induced fibril destabilization and molecular reorganization. The proposed framework may guide future experimental studies aimed at controlling protein aggregation through external physical fields, providing a theoretical basis for exploring new strategies to modulate amyloid structures.

## 5 Methods

### 5.1 System Preparation

Three A*β* fibril (2LMP, 2MXU, and 5OQV) were selected to investigate their structural responses to intense THz and mid-infrared pulses. As shown in Figure 1(a), 2LMP consists of 18 A*β*_40_ peptide chains arranged in a threefold symmetric structure, whereas 2MXU contains 12 A*β*_42_ chains with a different stacking arrangement. The 5OQV model comprises nine A*β*_42_ chains extracted from a parent structure containing two protofilaments. However, this nine-chain fragment was not assumed to exhibit strict twofold rotational symmetry. These models allow comparison of fibrils with different peptide compositions and packing arrangements.

All MD simulations were performed using GROMACS 2020.4 with the AMBER99SBILDN force field and the TIP4P water model. Each fibril was placed in a truncated octahedral simulation box, and Na^+^ or Cl*^−^* ions were added to neutralize the net charge. Energy minimization was performed using the steepest descent method to remove unfavorable atomic contacts. The Verlet cutoff scheme was used for nonbonded interactions, and long-range electrostatic interactions were calculated using the particle mesh Ewald (PME) method.

Following energy minimization, each system underwent 1 ns of NPT equilibration to adjust the system density and pressure. The leap-frog integration algorithm was used with positional restraints applied to the protein, while pressure was controlled using isotropic Parrinello–Rahman coupling. This was followed by 20 ns of NVT equilibration at 310 K, using the velocity-rescaling thermostat with separate temperature-coupling groups for the protein and solvent. The Verlet cutoff scheme and PME method were retained for nonbonded and long-range electrostatic interactions, respectively. Additional simulation parameters are provided in Table S1 of the Supporting Information.

These preparation and equilibration steps were used to obtain stable initial configurations and solvent environments. Configurations selected from the equilibrated NVT trajectories were then used as starting structures for NEMD simulations at different pulse frequencies and electric-field strengths.

### 5.2 Equilibrium and High-Temperature Control Simulations

To assess the structural stability of A*β* fibrils at different temperatures and provide reference data for the pulse-induced NEMD simulations, field-free control simulations (*E*_0_ = 0) were performed at 310 and 500 K. Both simulations were conducted in the NVT ensemble without pressure coupling.

For the 310 K control, the leap-frog integration algorithm was used with a time step of 1 fs for a total of 100 ns. The protein and solvent groups were separately coupled to 310 K using the Berendsen thermostat (*41*). Nonbonded interactions were handled using the Verlet neighbor-search scheme, with the neighbor list updated every 50 integration steps. Long-range electrostatic interactions were calculated using the PME method. This simulation served as a reference for fibril stability near physiological temperature.

For the high-temperature control, the target temperature was set to 500 K, with a time step of 2 fs and a total simulation time of 200 ns. All other simulation settings were identical to those of the 310 K control. This simulation was used to examine changes in secondary structure and hydrogen-bond networks under thermal perturbation and to compare them with pulse-induced structural changes.

These field-free simulations provided reference data for evaluating fibril responses under different conditions. However, the energy input and temperature histories were not matched between the high-temperature and pulse-driven simulations. Therefore, these comparisons were used to assess differences in structural evolution and intermolecular interactions, rather than to exclude possible thermal effects of the pulses. Detailed parameters are provided in Table S2 of the Supporting Information.

### 5.3 Pulsed-Field Nonequilibrium Molecular Dynamics Simulations

NEMD simulations were performed using GROMACS to investigate the structural responses of A*β* fibrils to pulsed electric fields over 0.5– 60 THz. A frequency interval of 0.5 THz was used, giving 120 nonzero frequency points. Each simulation lasted 50 ps with a time step of 0.2 fs. The applied field was a Gaussian-modulated cosine pulse, as illustrated in Figure 1b, with a peak field strength of 2 V*/*nm, a pulse center at 10 ps, and a Gaussian envelope standard deviation of 4 ps. Three independent simulations were performed at each frequency to assess reproducibility. This setup allowed comparison of picosecond pulse responses across different frequencies, including 50 THz in the amide I region.

The electric field was spatially uniform and polarized along the *x*-axis, with the *z*-axis representing the nominal propagation direction. Electromagnetic wave propagation was not explicitly simulated. The time-dependent electric field was defined as

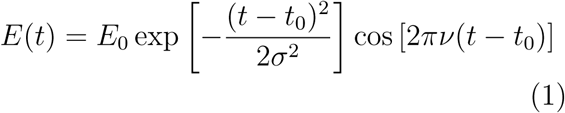

where *E*_0_ is the peak electric-field amplitude, varied from 0 to 2 V*/*nm; *σ* = 4 ps is the Gaussian envelope standard deviation; *t*_0_ = 10 ps is the pulse center; and *ν* is the carrier frequency, ranging from 0.5 to 60 THz. The phase is dimensionless when frequency and time are expressed in THz and ps, respectively.

For each fibril model (2LMP, 2MXU, and 5OQV), three configurations were selected from the end of the NVT equilibration trajectory as starting structures for independent simulations. Each NEMD trajectory was continued from an equilibrated checkpoint file. The Berendsen thermostat was applied only to water and ions at 310 K, while the protein was not directly coupled to a heat bath, allowing its temperature to respond to the applied field. All simulations were performed at constant volume without pressure coupling. Nonbonded interactions were treated using the Verlet cutoff scheme, with long-range electrostatics calculated by PME and van der Waals interactions handled using a simple cutoff. The neighbor list was updated every 10 fs. No bond constraints were applied to the protein to retain high-frequency vibrational degrees of freedom. Detailed simulation parameters are provided in Table S2 of the Supporting Information.

### 5.4 Calculation of Dipole Power Spectra and Absorption Spectrum

To compare protein dipole fluctuations and frequency-domain responses under different conditions, dipole power spectra and classical absorption proxies were calculated from independent 125 ps field-free MD trajectories. The integration time step was 1 fs, and coordinates were saved every 5 fs, yielding 25 001 frames per trajectory and a Nyquist frequency of 100 THz.

Protein dipole moments were calculated using the gmx dipoles tool in GROMACS 2020.4, with atomic charges taken from the simulation topology based on the AMBER99SBILDN force field. Following the molecular integrity treatment and center-of-mass reference convention for charged molecules, the total dipole vector was defined as

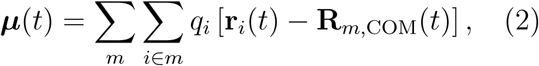

where *m* denotes a protein molecule defined in the topology, and *q_i_*, **r***_i_*(*t*), and **R***_m,_*_COM_(*t*) are the atomic charge, atomic position, and molecular center-of-mass position, respectively.

The time-averaged dipole vector was subtracted from each component, followed by Hann windowing:

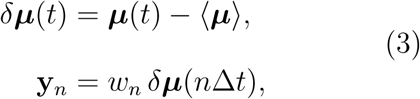

where *w_n_* is the Hann window and Δ*t* = 5 fs. The windowed linear vector autocorrelation function was estimated as

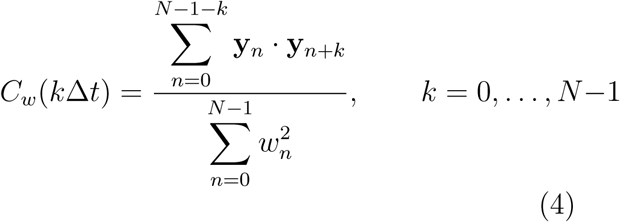

where *N* is the number of sampled frames. Sufficient zero-padding was applied to avoid circular correlation artifacts. Fluctuation amplitudes were preserved without normalization by *C_w_*(0).

The two-sided dipole power spectral density, *S_µ_*(*ν*), was calculated from the windowed series, and its nonnegative-frequency portion was retained. This is equivalent to Fourier transforming the corresponding autocorrelation function after full even extension. The classical absorption proxy was defined as

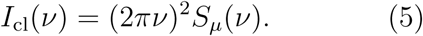

This quantity was used to compare the frequency-dependent spectral weight of protein dipole fluctuations. It excludes solvent and protein–solvent cross-correlation contributions and does not represent the absolute absorption coefficient of the system.

Spectra were calculated separately for each trajectory. For conditions with independent replicates, equally weighted mean spectra and sample standard deviations across replicates were calculated on a common frequency grid. The number of valid replicates is specified in the corresponding figure captions. Spectra are primarily presented over 0–60 THz. For the comparison of 2LMP spectra before and after pulse exposure, all curves were normalized by the maximum intensity of the unnormalized pre-pulse 2LMP spectrum within this range, preserving relative intensity differences between conditions.

### 5.5 Bond-Angle Analysis

To characterize local conformational changes in the protein, bond angles were calculated for three sequentially bonded atoms. For atoms with positions **r***_i_*, **r***_j_*, and **r***_k_*, the bond angle centered at atom *j* was defined as

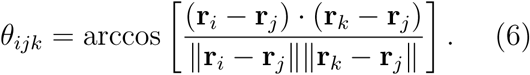

The angle *θ_ijk_*ranges from 0 to *π* radians. Its time evolution and distribution were analyzed to characterize pulse-induced local conformational changes.

---

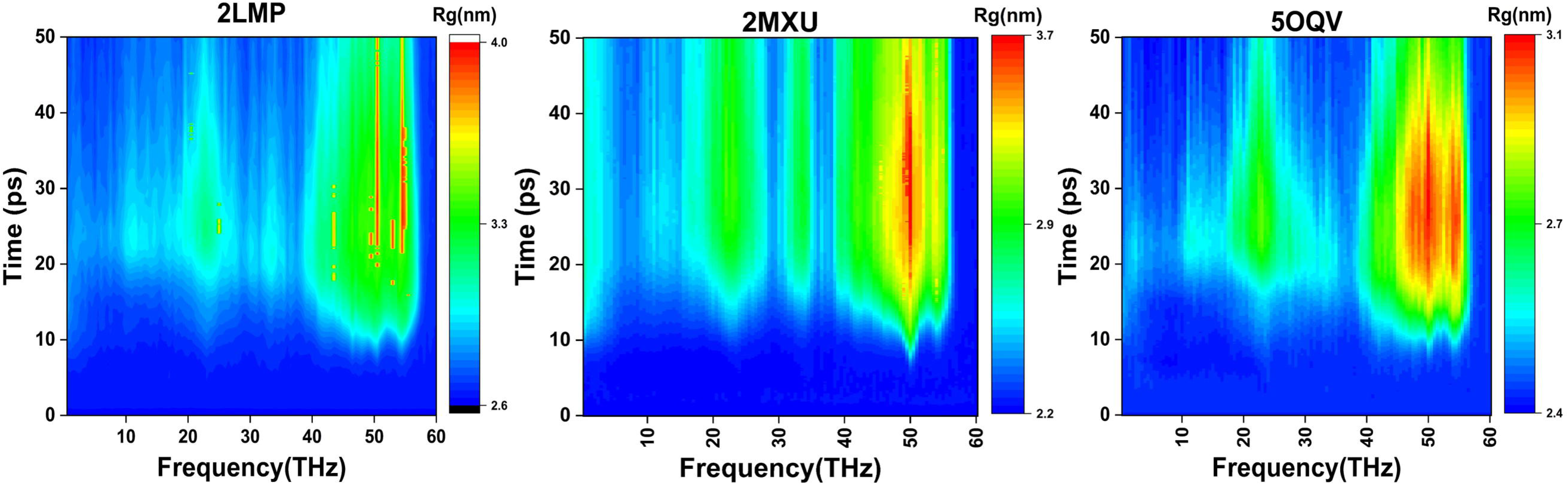

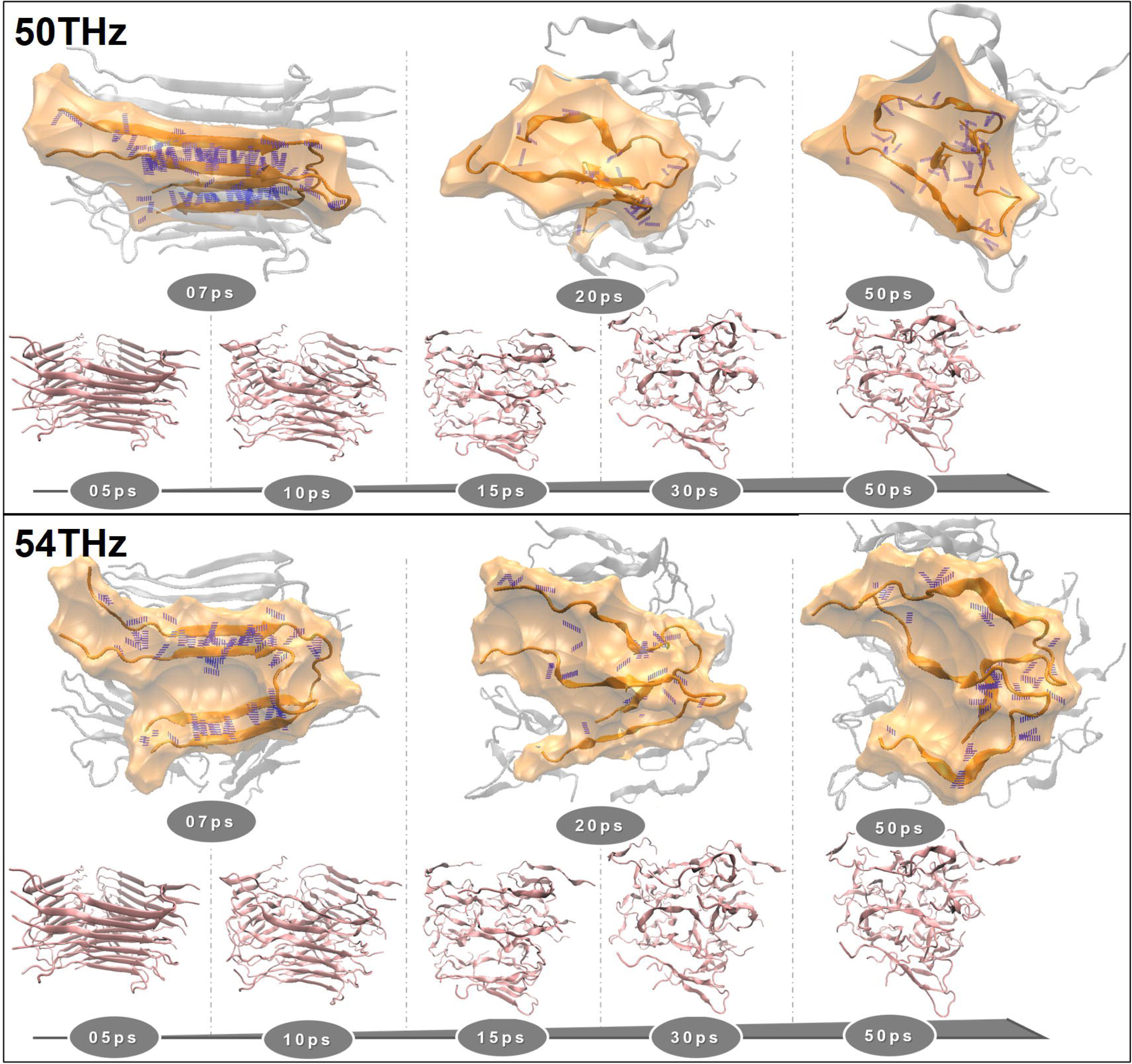

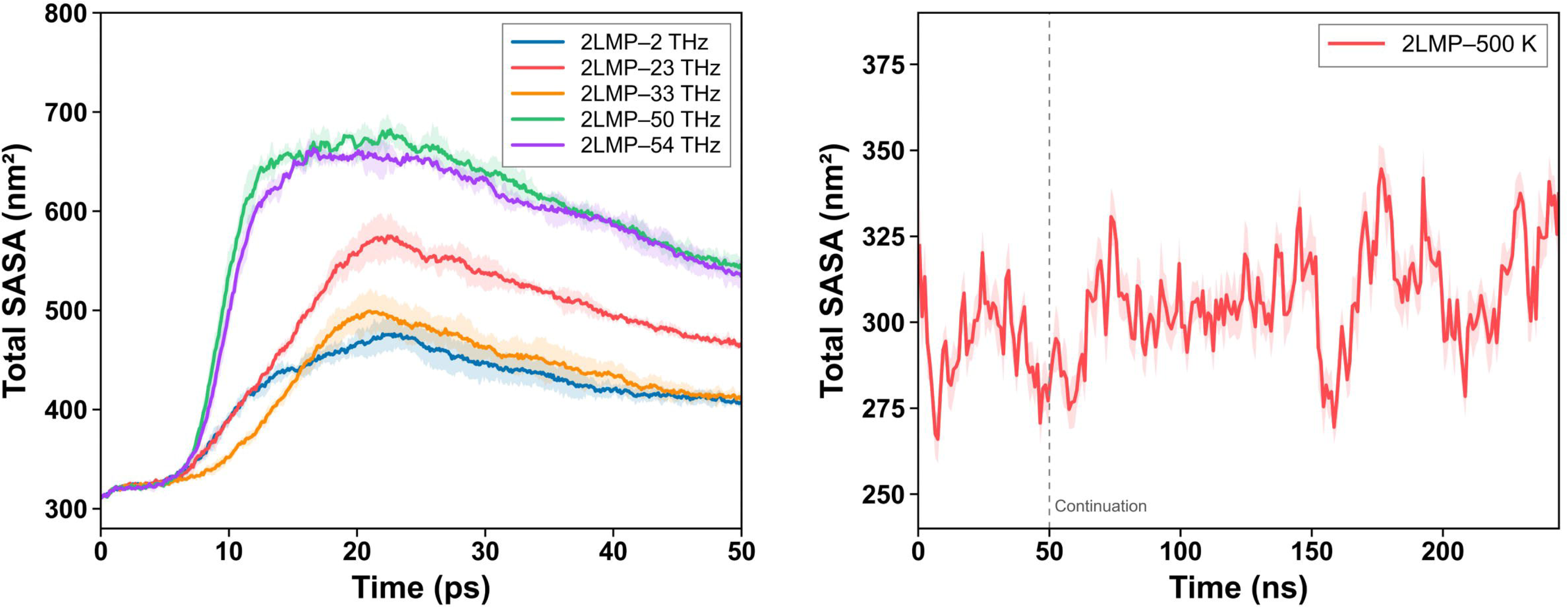

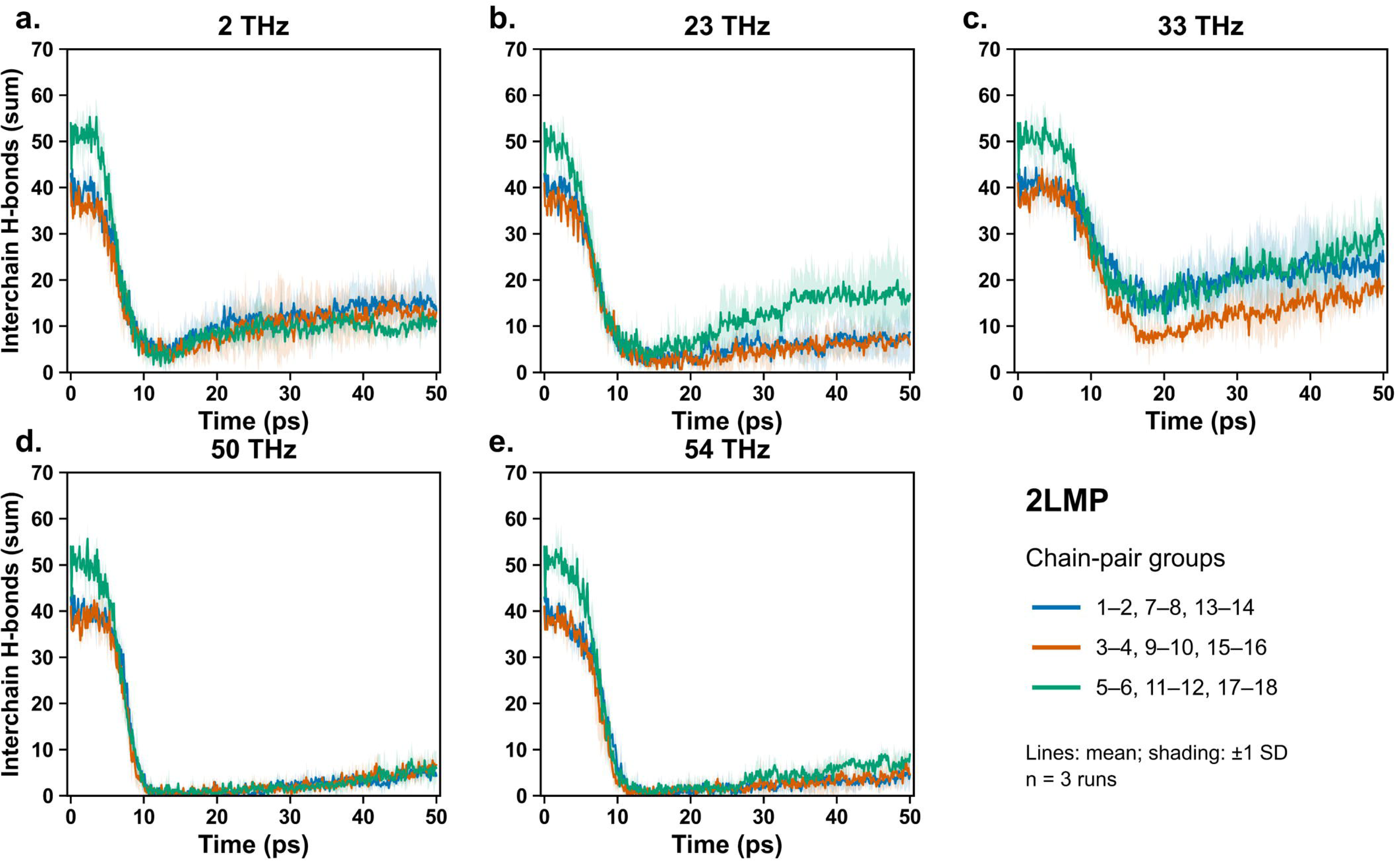

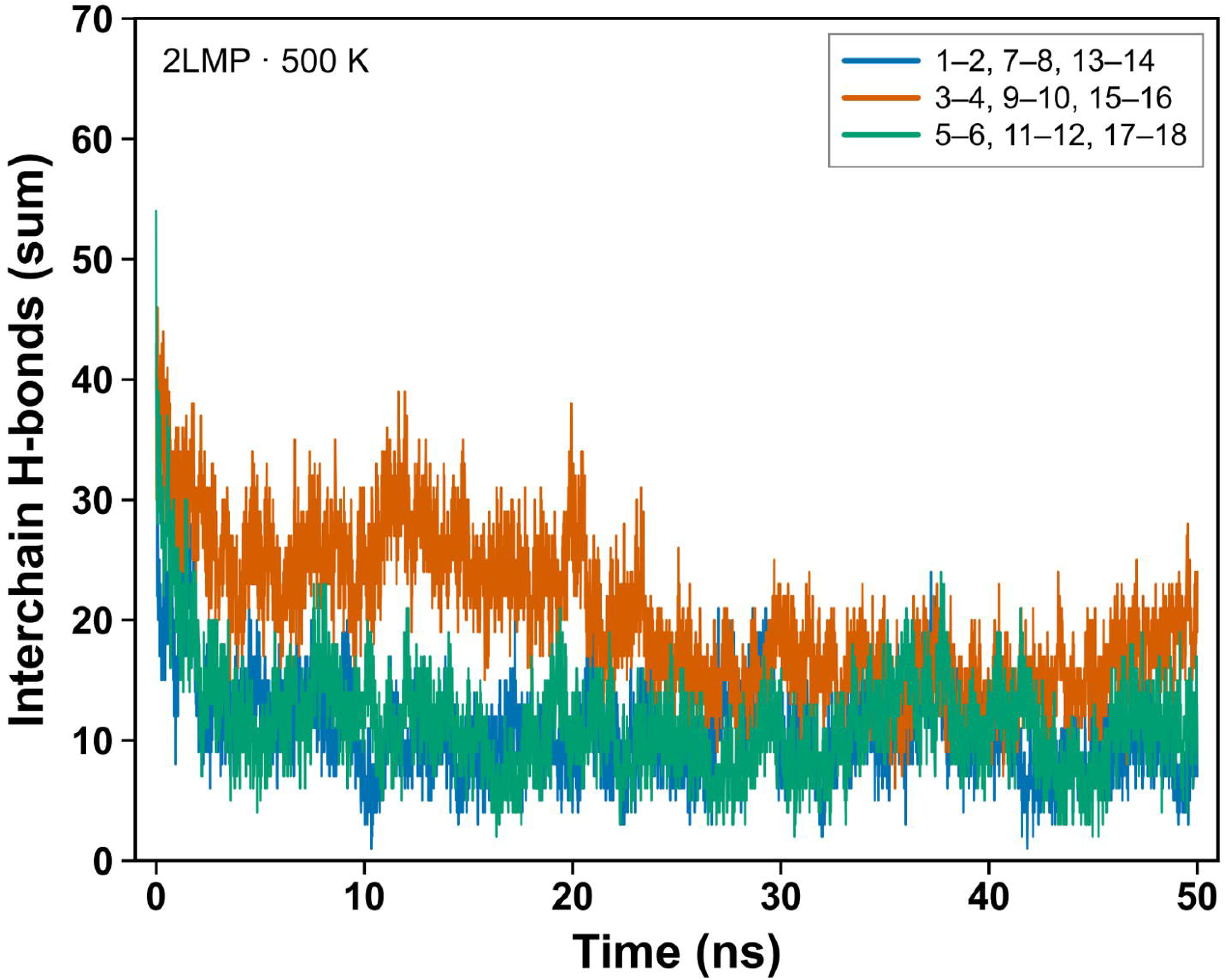

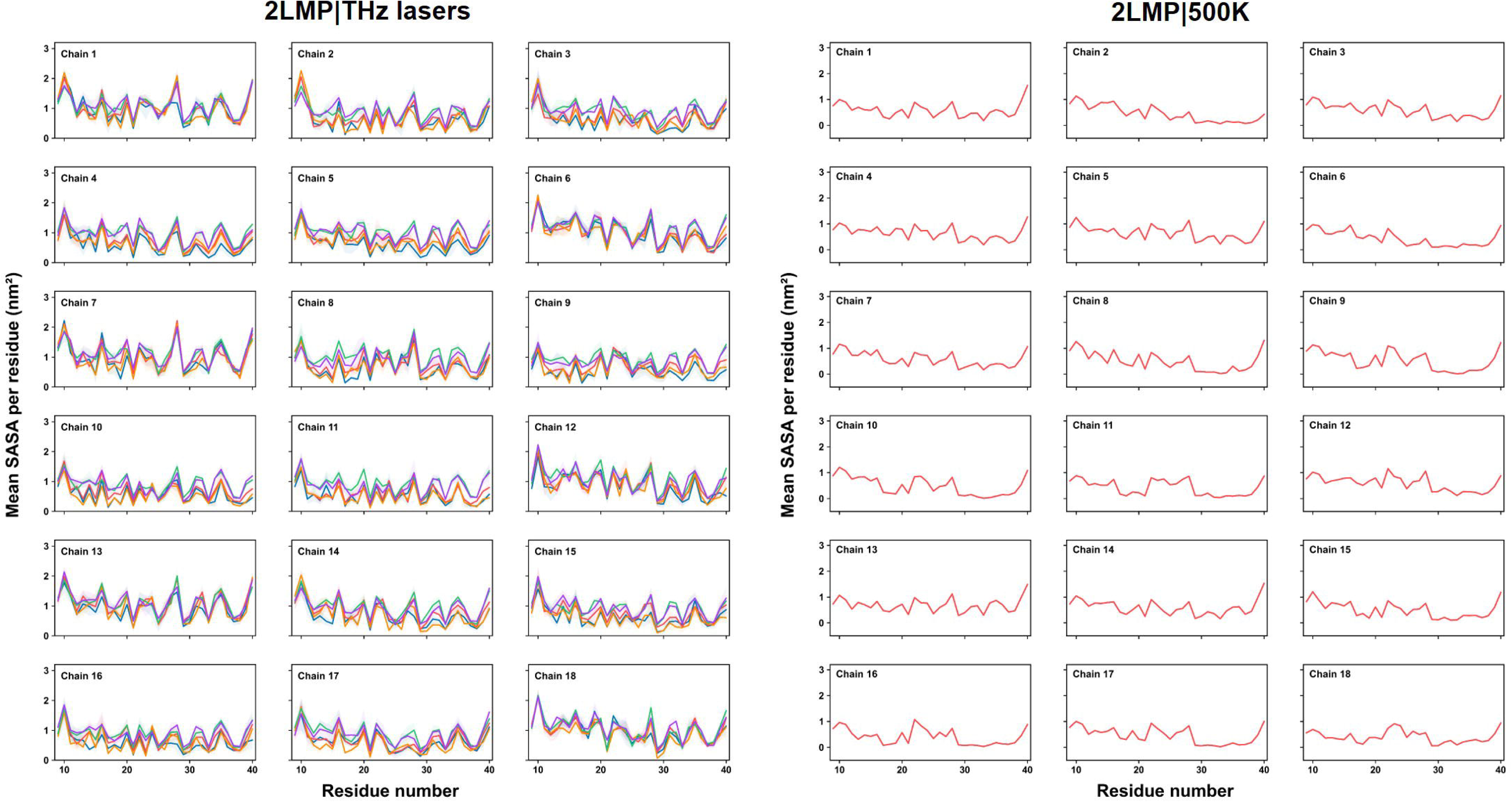

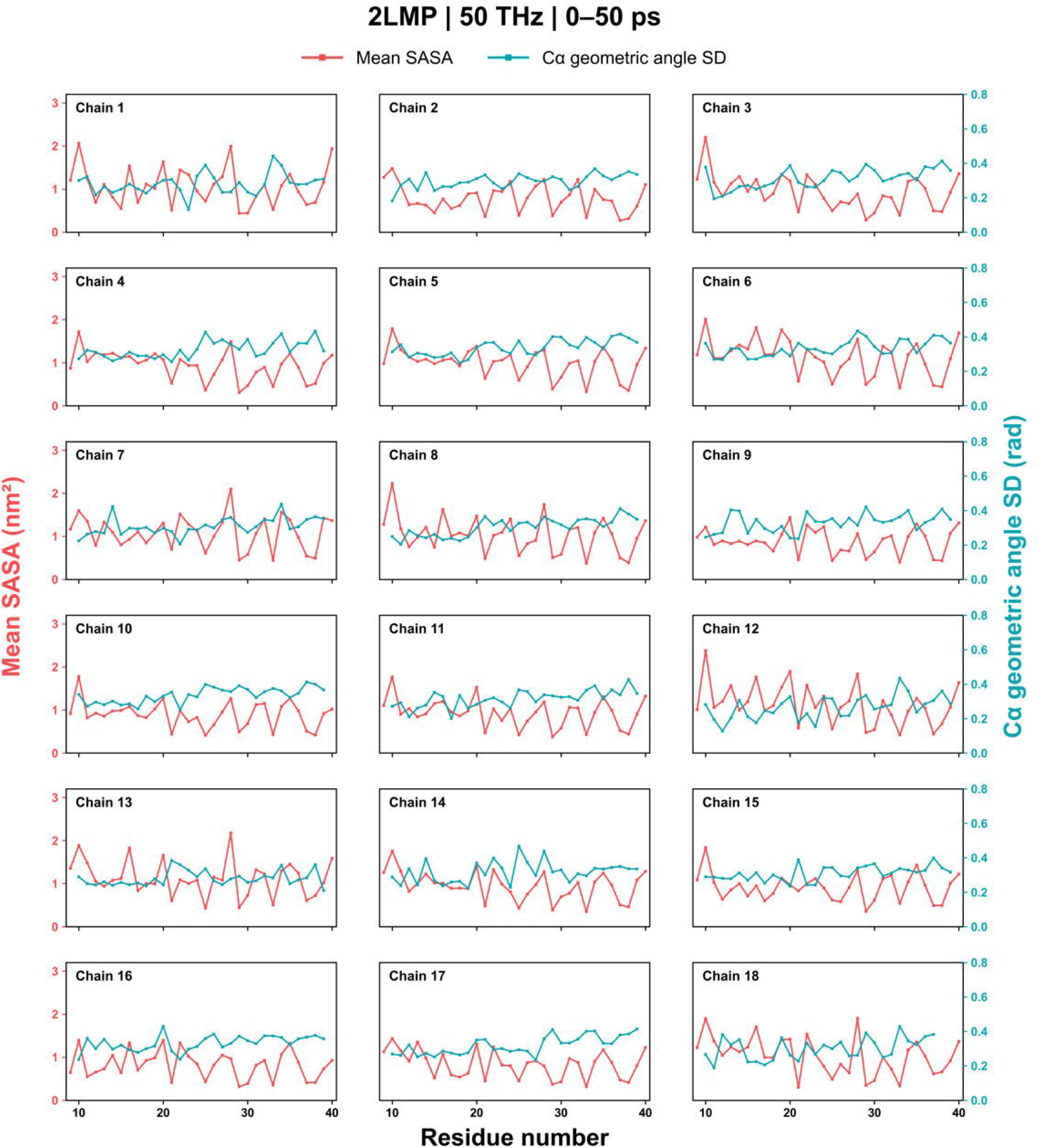

## References

1. Friedland, R. P., and Haribabu, B. (2023) Neurodegenerative diseases: from gut-brain axis to brain microbiome. Frontiers in Aging Neuroscience 15, 1171955.

2. Chen, H., Zeng, Y., Wang, D., Li, Y., Xing, J., Zeng, Y., Liu, Z., Zhou, X., and Fan, H. (2024) Neuroinflammation of Microglial Regulation in Alzheimer’s Disease: Therapeutic Approaches. Molecules 29, 1478.

3. Bai, R., Guo, J., Ye, X.-Y., Xie, Y., and Xie, T. (2022) Oxidative stress: The core pathogenesis and mechanism of Alzheimer’s disease. Ageing Research Reviews 77, 101619.

4. Ju, Y., and Tam, K. Y. (2022) Pathological mechanisms and therapeutic strategies for Alzheimer’s disease. Neural Regeneration Research 17, 543–549.

5. Haut, F., Argyrousi, E. K., and Arancio, O. (2024) Re-Arranging the Puzzle between the Amyloid-Beta and Tau Pathology: An APP-Centric Approach. International Journal of Molecular Sciences 25, 259.

6. Du, Z., Li, M., Ren, J., and Qu, X. (2021) Current Strategies for Modulating A*β* Aggregation with Multifunctional Agents. Accounts of Chemical Research 54, 2172–2184.

7. Sakono, M., and Zako, T. (2010) Amyloid oligomers: formation and toxicity of A*β* oligomers. The FEBS Journal 277, 1348–1358.

8. Ferreira, S. T., Lourenco, M. V., Oliveira, M. M., and De Felice, F. G. (2015) Soluble amyloid-*β* oligomers as synaptotoxins leading to cognitive impairment in Alzheimer’s disease. Frontiers in Cellular Neuroscience 9, 191.

9. Kaku, T., Ikebukuro, K., and Tsukakoshi, K. (2024) Structure of cytotoxic amyloid oligomers generated during disaggregation. The Journal of Biochemistry 175, 575–585.

10. Jurcău, M. C., Andronie-Cioara, F. L., Jurcău, A., Marcu, F., Ţiţ, D. M., Paşcalău, N., and Nistor-Cseppentő, D. C. (2022) The link between oxidative stress, mitochondrial dysfunction and neuroinflammation in the pathophysiology of Alzheimer’s disease: therapeutic implications and future perspectives. Antioxidants 11, 2167.

11. Cascella, R., and Cecchi, C. (2021) Calcium dyshomeostasis in Alzheimer’s disease pathogenesis. International Journal of Molecular Sciences 22, 4914.

12. Guan, P.-P., Cao, L.-L., Yang, Y., and Wang, P. (2021) Calcium Ions Aggravate Alzheimer’s Disease Through the Aberrant Activation of Neuronal Networks, Leading to Synaptic and Cognitive Deficits. Frontiers in Molecular Neuroscience 14, 757515.

13. Almeida, Z. L., and Brito, R. M. M. (2020) Structure and aggregation mechanisms in amyloids. Molecules 25, 1195.

14. Chen, G.-f., Xu, T.-h., Yan, Y., Zhou, Y.-r., Jiang, Y., Melcher, K., and Xu, H. E. (2017) Amyloid beta: structure, biology and structure-based therapeutic development. Acta Pharmacologica Sinica 38, 1205–1235.

15. Ghosh, U., Thurber, K. R., Yau, W.-M., and Tycko, R. (2021) Molecular structure of a prevalent amyloid-beta fibril polymorph from Alzheimer’s disease brain tissue. Proceedings of the National Academy of Sciences 118, e2023089118.

16. Cario, A., and Berger, C. L. (2023) Tau, microtubule dynamics, and axonal transport: New paradigms for neurodegenerative disease. BioEssays 45, 2200138.

17. Mercken, M., Fischer, I., Kosik, K. S., and Nixon, R. A. (1995) Three distinct axonal transport rates for tau, tubulin, and other microtubule-associated proteins: evidence for dynamic interactions of tau with microtubules in vivo. Journal of Neuroscience 15, 8259–8267.

18. Sarkar, S. (2018) Neurofibrillary tangles mediated human neuronal tauopathies: insights from fly models. Journal of Genetics 97, 783–793.

19. Wu, M., Zhang, M., Yin, X., Chen, K., Hu, Z., Zhou, Q., Cao, X., Chen, Z., and Liu, D. (2021) The role of pathological tau in synaptic dysfunction in Alzheimer’s diseases. Translational Neurodegeneration 10, 45.

20. Malpas, C. B., Sharmin, S., and Kalincik, T. (2021) The histopathological staging of tau, but not amyloid, corresponds to antemortem cognitive status, dementia stage, functional abilities and neuropsychiatric symptoms. International Journal of Neuroscience 131, 800–809.

21. Haddad, H. W., Malone, G. W., Comardelle, N. J., Degueure, A. E., Poliwoda, S., Kaye, R. J., Murnane, K. S., Kaye, A. M., and Kaye, A. D. (2022) Aduhelm, a novel anti-amyloid monoclonal antibody, for the treatment of Alzheimer’s Disease: A comprehensive review. Health Psychology Research 10, 37023.

22. Wu, K., Kwon, S. H., Zhou, X., Fuller, C., Wang, X., Vadgama, J., and Wu, Y. (2024) Overcoming Challenges in Small-Molecule Drug Bioavailability: A Review of Key Factors and Approaches. International Journal of Molecular Sciences 25, 13121.

23. Rouhi, N., Chakeri, Z., Nejad, B. G., Rahimzadegan, M., Khezri, M. R., Kamali, H., and Nosrati, R. (2024) A comprehensive review of advanced focused ultrasound (FUS) microbubbles-mediated treatment of Alzheimer’s disease. Heliyon 10.

24. Nilsson, P., Iwata, N., Muramatsu, S.-i., Tjernberg, L. O., Winblad, B., and Saido, T. C. (2010) Gene therapy in Alzheimer’s disease – potential for disease modification. Journal of Cellular and Molecular Medicine 14, 741–757.

25. Zhang, Y., Chen, H., Li, R., Sterling, K., and Song, W. (2023) Amyloid *β*-based therapy for Alzheimer’s disease: challenges, successes and future. Signal Transduction and Targeted Therapy 8, 248.

26. Ramundo Orlando, A., and Gallerano, G. P. (2009) Terahertz radiation effects and biological applications. Journal of Infrared, Millimeter, and Terahertz Waves 30, 1308–1318.

27. Nikitkina, A. I., Bikmulina, P. Y., Gafarova, E. R., Kosheleva, N. V., Efremov, Y. M., Bezrukov, E. A., Butnaru, D. V., Dolganova, I. N., Chernomyrdin, N. V., Cherkasova, O. P., Gavdush, A. A., and Timashev, P. S. (2021) Terahertz radiation and the skin: a review. Journal of Biomedical Optics 26, 043005.

28. Mancini, T., Mosetti, R., Marcelli, A., Petrarca, M., Lupi, S., and D’Arco, A. (2022) Terahertz Spectroscopic Analysis in Protein Dynamics: Current Status. Radiation 2, 100–123.

29. Pezzotti, S., Chen, W., Novelli, F., Yu, X., Hoberg, C., and Havenith, M. (2025) Terahertz calorimetry spotlights the role of water in biological processes. Nature Reviews Chemistry 9, 481–494.

30. Sugiyama, J.-i., Tokunaga, Y., Hishida, M., Tanaka, M., Takeuchi, K., Satoh, D., and Imashimizu, M. (2023) Nonthermal acceleration of protein hydration by sub-terahertz irradiation. Nature Communications 14, 2825.

31. Govorun, V. M., Tretiakov, V. E., Tulyakov, N. N., Fleurov, V. B., Demin, A. I., Volkov, A. Y., Batanov, V. A., and Kapitanov, A. B. (1991) Far-infrared radiation effect on the structure and properties of proteins. International Journal of Infrared and Millimeter Waves 12, 1469–1474.

32. Homenko, A., Kapilevich, B., Kornstein, R., and Firer, M. (2009) Effects of 100 GHz radiation on alkaline phosphatase activity and antigen–antibody interaction. Bioelectromagnetics 30, 167–175.

33. Zhang, J., Chen, Y., Zhao, Y., Wang, P., Ding, H., Liu, C., Lyu, J., and Le, W. (2024) Terahertz Irradiation Improves Cognitive Impairments and Attenuates Alzheimer’s Neuropathology in the APPSWE/PS1DE9 Mouse: A Novel Therapeutic Intervention for Alzheimer’s Disease. Neuroscience Bulletin 40, 857–871.

34. Acbas, G., Niessen, K. A., Snell, E. H., and Markelz, A. (2014) Optical measurements of long-range protein vibrations. Nature Communications 5, 3076.

35. Peng, W., Zhu, Z., Lou, J., Chen, K., Wu, Y., and Chang, C. (2023) High-frequency terahertz waves disrupt Alzheimer’s *β*-amyloid fibril formation. eLight 3, 18.

36. Pu, Z., Wu, Y., Zhu, Z., Zhao, H., and Cui, D. (2025) A new horizon for neuroscience: terahertz biotechnology in brain research. Neural Regeneration Research 20, 309–325.

37. Fellows, A. P., Casford, M. T. L., and Davies, P. B. (2020) Spectral Analysis and Deconvolution of the Amide I Band of Proteins Presenting with High-Frequency Noise and Baseline Shifts. Applied Spectroscopy 74, 597–615.

38. Kawasaki, T., Tsukiyama, K., and Nguyen, P. H. (2023) Disassembly of Amyloid Fibril with Infrared Free Electron Laser. International Journal of Molecular Sciences 24, 3686.

39. Hoang Viet, M., Derreumaux, P., Li, M. S., Roland, C., Sagui, C., and Nguyen, P. H. (2015) Picosecond dissociation of amyloid fibrils with infrared laser: A nonequilibrium simulation study. The Journal of Chemical Physics 143, 155101.

40. Kawasaki, T., Man, V. H., Sugimoto, Y., Sugiyama, N., Yamamoto, H., Tsukiyama, K., Wang, J., Derreumaux, P., and Nguyen, P. H. (2020) Infrared Laser-Induced Amyloid Fibril Dissociation: A Joint Experimental/Theoretical Study on the GNNQQNY Peptide. The Journal of Physical Chemistry B 124, 6266–6277.

41. Berendsen, H. J. C., Postma, J. P. M., van Gunsteren, W. F., DiNola, A., and Haak, J. R. (1984) Molecular dynamics with coupling to an external bath. The Journal of Chemical Physics 81, 3684–3690.

