## Supplemental Information for "Frequency-Dependent Structural Responses of Amyloid- Fibrils to Terahertz Pulses: A Nonequilibrium Molecular Dynamics Study"

This Supplementary Information provides the simulation settings and additional structural analyses accompanying the six main-text figures. Tables S1 and S2 summarize the production protocols. Figures S1 and S2 address global size and selected conformations; Figures S3–S7 describe solvent exposure, interchain hydrogen bonds, and local backbone geometry. Figures S8 and S9 are reserved for the spectral comparison and the two-threshold classification method, respectively.

#### S1 Simulation systems and production protocols

The three fibril models were derived from PDB entries 2LMP, 2MXU, and 5OQV[1–3]. The simulated models contain 18 chains of A$\beta$40, 12 chains of A$\beta$42, and a nine-chain A$\beta$42 fibril fragment, respectively. The parent sequence length does not specify the number of residues retained in the simulation. Likewise, the organization of the parent 5OQV fibril does not establish an exact twofold symmetry for the selected nine-chain fragment.

The systems were solvated with TIP4P water[4] and neutralized with counterions. System preparation included energy minimization, $1\ \mathrm{ns}$ of NPT equilibration, and $20\ \mathrm{ns}$ of NVT equilibration at $310\ \mathrm{K}$, as described in the main text. Production simulations used GROMACS 2020.4[5], the AMBER-99SB-ILDN protein force field[6], particle mesh Ewald electrostatics[7], and Berendsen temperature coupling[8] with the protocol-specific settings in Table S2.

**Table S1.** Fibril systems and common production molecular dynamics settings.

| Parameter | Specification |
|---|---|
| Fibril structures (PDB identifiers) | 2LMP, 2MXU, and 5OQV |
| Number of replicates | Three per investigated system–condition combination (protocol specification) |
| Simulation software | GROMACS 2020.4 |
| Protein force field | AMBER-99SB-ILDN |
| Integration algorithm | Leap-frog molecular dynamics |
| Periodic boundary conditions | Applied along the $x$, $y$, and $z$ directions |
| Pressure coupling | None; fixed simulation box |
| Neighbor-search scheme | Verlet cutoff scheme |
| Configured neighbor-list update interval | 50 integration steps |
| Long-range electrostatics | Particle mesh Ewald (PME) |
| Real-space electrostatic cutoff | $1.2\ \mathrm{nm}$ |
| PME interpolation order | Fourth order |
| PME Fourier grid spacing | $0.16\ \mathrm{nm}$ |
| Temperature-coupling algorithm | Berendsen |
| Production-run initialization | Continuation enabled; no new velocities generated |
| Compressed trajectory output | Protein coordinates only |

Protocol-specific integration time steps, simulation durations, temperature-coupling settings, Lennard–Jones treatments, and electric-field parameters are summarized in Table S2. These settings apply to production runs, not to the preceding NPT equilibration. The replicate count describes the simulation protocol; the number of runs actually included in an individual plot is stated in its caption where available.

**Table S2.** Production simulation parameters for the 310 K control, 500 K heating control, and THz frequency scan.

| Parameter | 310 K control | 500 K heating control | THz frequency scan |
|---|---|---|---|
| Integration time step | 1 fs | 2 fs | 0.2 fs |
| Number of integration steps | $1.0 \times 10^8$ | $1.0 \times 10^8$ | $2.5 \times 10^5$ |
| Configured duration[a] | 100 ns | 200 ns | 50 ps |
| Temperature-coupling groups | Protein; non-protein | Protein; non-protein | Protein; water and ions |
| Protein reference temperature | 310 K | 500 K | Not coupled[b] |
| Non-protein bath temperature | 310 K | 500 K | 310 K |
| Protein coupling time constant | 0.1 ps | 0.1 ps | Disabled |
| Non-protein coupling time constant | 0.1 ps | 0.1 ps | 0.1 ps |
| Lennard–Jones cutoff | 1.1 nm | 1.1 nm | 1.2 nm |
| Lennard–Jones modifier[c] | Force switch | Force switch | Potential shift |
| Force-switching interval | 1.0–1.1 nm | 1.0–1.1 nm | — |
| Energy, log, and compressed trajectory output interval | 10 ps | 10 ps | 0.1 ps |
| Applied electric field | None | None | Gaussian-envelope oscillating field |
| Field direction | — | — | $x$ axis |
| Field amplitude, $E_0$ | 0 | 0 | $2\ \mathrm{V\,nm^{-1}}$ |
| Frequency range | — | — | 0.5–60 THz |
| Frequency increment | — | — | 0.5 THz |
| Number of scanned frequencies | — | — | 120 |
| Pulse-center parameter, $t_0$ | — | — | 10 ps |
| Gaussian-envelope width parameter, $\sigma$ | — | — | 4 ps |

[a] Configured durations are the products of the integration time step and the specified number of steps. Duplicate specifications of a heating protocol are not added to obtain a longer trajectory. For a continued run, the elapsed trajectory time and the analyzed window must be established from its run records.

[b] Protein temperature coupling was disabled in the THz simulations by setting `tau_t = -1`. The corresponding protein `ref_t` entry therefore does not impose a target temperature. Water and ions were coupled to a 310 K bath with a time constant of 0.1 ps.

[c] Force switching is specified for the control protocols. The THz protocol uses the GROMACS 2020.4 default potential-shift treatment when the van der Waals modifier is left unspecified[9].

The THz column describes the frequency scan at $E_0 = 2\ \mathrm{V\,nm^{-1}}$. The Gaussian-envelope parameter $\sigma$ is distinct from the full width at half maximum.

#### S2 Pulse definition and reporting conventions

The applied field is spatially uniform and polarized along the simulation $x$ axis:

$$\mathbf{E}(t) = E_0 \exp\left[-\frac{(t-t_0)^2}{2\sigma^2}\right] \cos[2\pi\nu(t-t_0)]\,\hat{\mathbf{x}}. \tag{S1}$$

Here $t_0 = 10$ ps and $\sigma = 4$ ps is the standard deviation of the field envelope, not its full width at half maximum. When $t$ is expressed in ps and the ordinary frequency $\nu$ in THz, the phase in Eq. (S1) is dimensionless. The GROMACS field parameter uses angular frequency, $\omega = 2\pi\nu$[9]. The optical propagation direction shown in schematics is not a spatially propagating wave in the simulation.

The frequency scan comprises 120 nonzero frequencies from $0.5$ to $60$ THz in increments of $0.5$ THz at $E_0 = 2\ \mathrm{V\,nm^{-1}}$. Each trajectory spans $50$ ps with a $0.2$ fs integration time step. The displayed label $33$ THz denotes the previously confirmed input frequency $33.5$ THz; it does not denote an additional simulation at $33.0$ THz. The representative main-text labels are 2, 23, 33, and $50$ THz, with $54$ THz retained for supplementary comparison. A field-free reference has $E_0 = 0$; setting only $\nu = 0$ does not remove a nonzero Gaussian field.

In the pulse protocol, water and ions are coupled to a $310$ K bath with a $0.1$ ps time constant. Protein temperature coupling is disabled. This differs from both temperature-control protocols, in which protein and non-protein groups are thermostatted. The $500$ K trajectory is a high-temperature structural reference. Its temperature and energy history are not matched to those of the pulse trajectories.

#### S3 Structural observables and statistical interpretation

##### S3.1 Global size and hydrogen-bond groups

The radius of gyration, $R_g$, describes the spatial extent of the selected protein atoms. For a mass-weighted selection it is defined by

$$R_g(t) = \left[\frac{\sum_i m_i |\mathbf{r}_i(t) - \mathbf{r}_{\mathrm{COM}}(t)|^2}{\sum_i m_i}\right]^{1/2}. \tag{S2}$$

The maps in Figure S1 complement the secondary-structure analysis. A change in overall size alone does not establish release of individual peptide chains.

For Figures S4 and S5, the plotted chain-pair groups are $G_1 = \{(1,2),(7,8),(13,14)\}$, $G_2 = \{(3,4),(9,10),(15,16)\}$, and $G_3 = \{(5,6),(11,12),(17,18)\}$. The group observable is the sum of the interchain hydrogen-bond counts over the three listed pairs,

$$N_{G_k}(t) = \sum_{(a,b)\in G_k} N_{a,b}(t). \tag{S3}$$

These groups sample selected interfaces; their sum is not automatically the count of all interchain hydrogen bonds in the fibril. For the full protein-protein hydrogen-bond population, a consistent partition instead satisfies $N_{\text{total}} = N_{\text{intra}} + N_{\text{inter}}$. Recovery of a count does not by itself demonstrate recovery of the original donor–acceptor pairs.

#### S3.2 Solvent exposure and local geometry

Total solvent-accessible surface area (SASA) is reported in $\mathrm{nm}^2$ in Figure S3. Residue-resolved time averages are shown for each of the 18 chains in Figure S6. These averages describe exposure over the stated observation window; post-excitation averages are not measures of initial burial. To classify residues by their initial environment, their SASA must instead be evaluated before excitation with the surrounding fibril included.

Local C$\alpha$ geometry requires three consecutive residues within the same peptide chain. With $\mathbf{t}_i = (\mathbf{r}_{i+1} - \mathbf{r}_i)/|\mathbf{r}_{i+1} - \mathbf{r}_i|$, the turning angle is

$$\kappa_i = \arccos(\mathbf{t}_{i-1} \cdot \mathbf{t}_i), \qquad \theta_i = \pi - \kappa_i, \tag{S4}$$

where $\theta_i$ is the corresponding three-point internal angle. Both describe coarse-grained backbone geometry rather than a covalent bond angle between three directly bonded atoms. A pseudodihedral requires four consecutive C$\alpha$ atoms. Cross-chain triples, missing-residue breaks, and undefined terminal positions must be excluded.

Figure S7 compares residue-resolved mean SASA with the temporal standard deviation of the C$\alpha$ geometric angle over $0$–$50$ ps. This dispersion includes both conformational transitions and shorter-time fluctuations. It is not a direct measure of vibrational energy.

#### S3.3 Replicates and time windows

Where a plot explicitly reports three runs, each run is the statistical unit. For a time-resolved observable $X_r(t)$ from $R = 3$ runs, the mean and sample standard deviation across runs are

$$\overline{X}(t) = \frac{1}{R}\sum_{r=1}^{R} X_r(t), \qquad s_X(t) = \left[\frac{1}{R-1}\sum_{r=1}^{R}\big(X_r(t) - \overline{X}(t)\big)^2\right]^{1/2}. \tag{S5}$$

For a residue-resolved time average, averaging within each run precedes summarizing the resulting run-level values. The shaded bands in Figure S4 and the pulse panel of Figure S6 are identified in the supplied plots as mean $\pm$ one standard deviation across three runs. In contrast, the high-temperature panel of Figure S3 reports means and temporal standard deviations within $1\ \mathrm{ns}$ bins of a single trajectory. These temporal fluctuations are not uncertainties from independent repeats. The high-temperature panel of Figure S6 likewise identifies one continuous run.

### S4 Supplementary figures

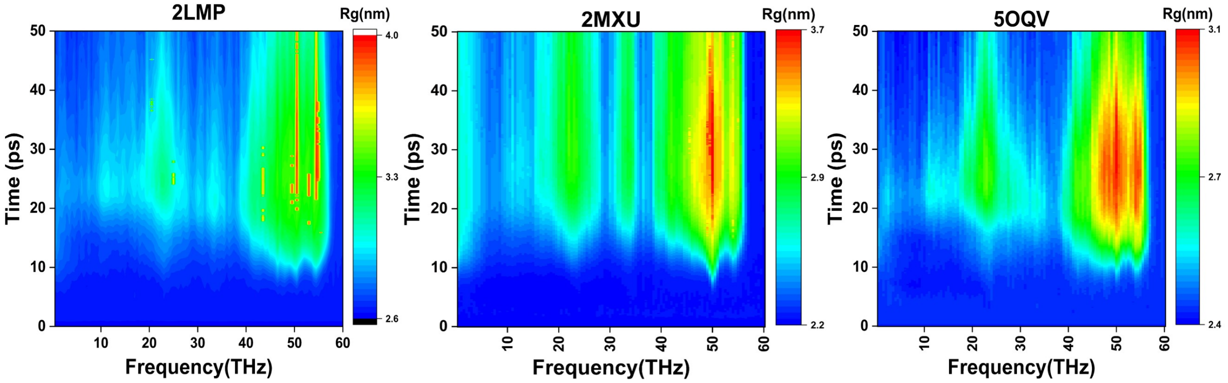


**Figure S1.** Frequency-dependent evolution of the radius of gyration of the three fibril models. The panels correspond to 2LMP, 2MXU, and 5OQV, as labeled. The protocol scans $0.5$–$60$ THz at $E_0 = 2\ \mathrm{V\,nm^{-1}}$ with $x$ polarization over $0$–$50$ ps. Color denotes $R_g$ in nm; each panel has its own color scale. The maps describe changes in overall protein size and complement the main-text secondary-structure maps.

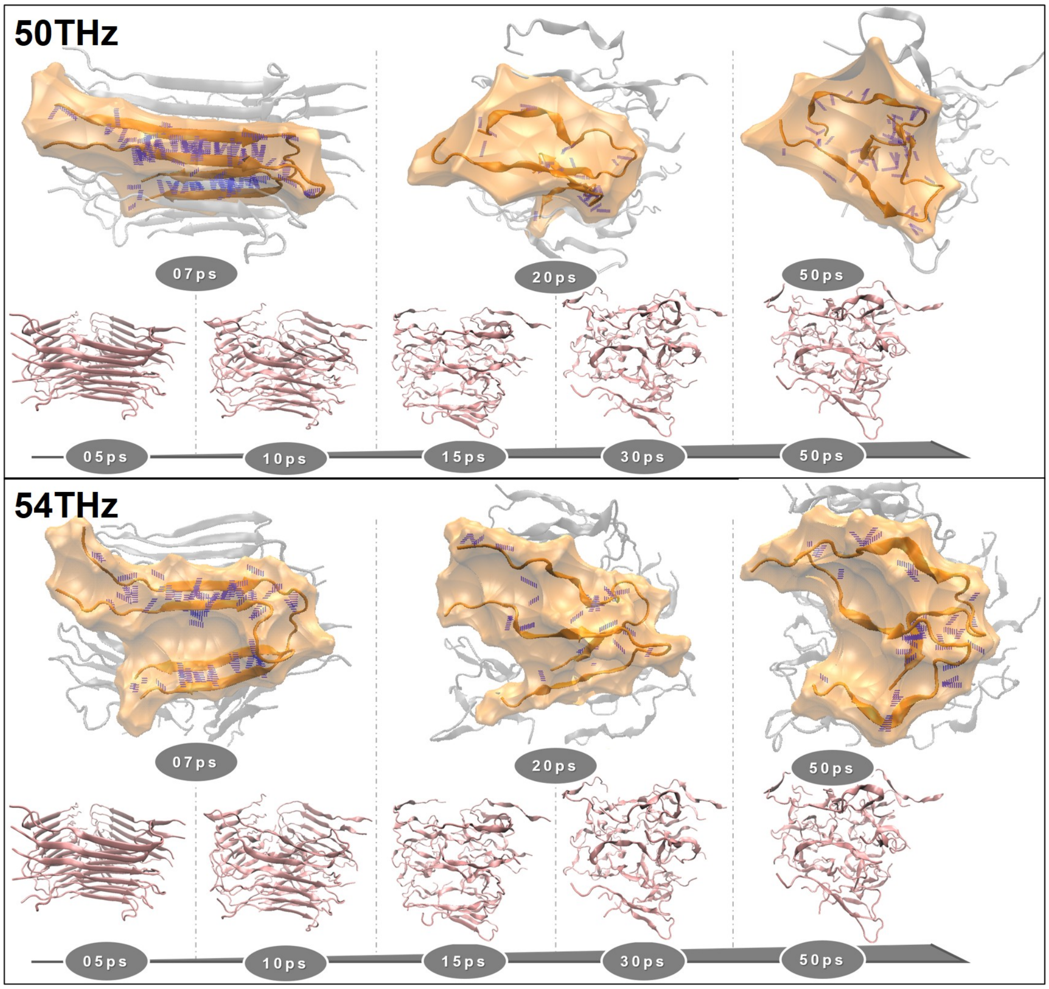


**Figure S2.** Selected conformational views for the 50 and 54 THz 2LMP pulse conditions. The upper and lower comparison blocks are labeled 50 and 54 THz, respectively, at $E_0 = 2\ \mathrm{V\,nm^{-1}}$ with $x$ polarization. Local views highlight chains 1 and 2 at 7, 20, and 50 ps; the whole-fibril timelines are labeled 5, 10, 15, 30, and 50 ps. Highlighted chains are orange, surrounding chains are gray, and the displayed hydrogen bonds are purple. These selected views provide structural context and do not establish identical vibrational modes or long-time irreversibility.

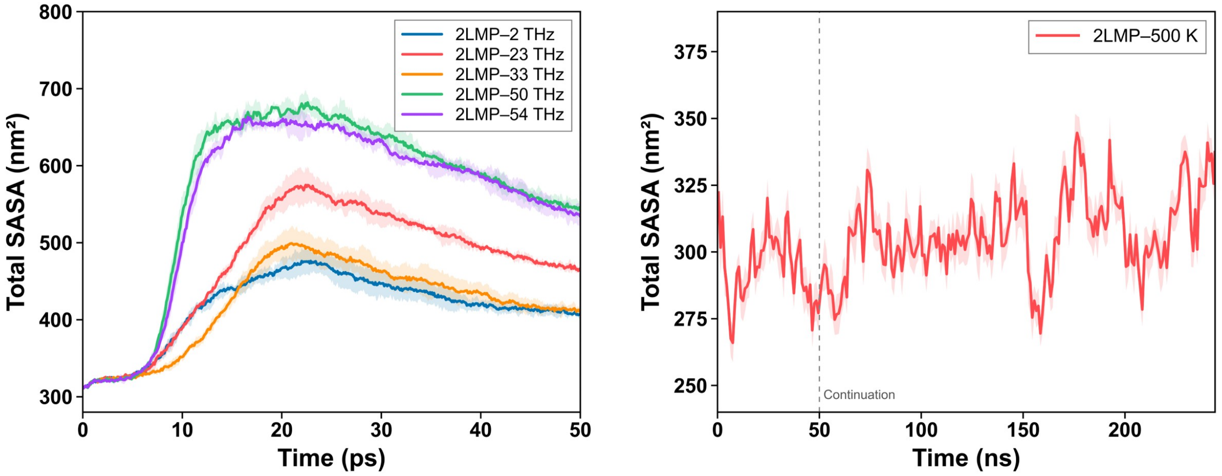


**Figure S3.** Time evolution of total SASA for 2LMP under pulse excitation and heating. Left: the 2, 23, 33, 50, and $54$ $\mathrm{THz}$ pulse conditions over $0$–$50$ $\mathrm{ps}$, shown in blue, red, orange, green, and purple. The peak field is $2$ $\mathrm{V\,nm^{-1}}$ and is polarized along $x$. Right: the $500$ $\mathrm{K}$ field-free trace with time in ns. Its line and shaded region denote the mean and temporal standard deviation within $1$ $\mathrm{ns}$ bins, as labeled in the supplied plot. The dashed line marks a continuation boundary near $50$ $\mathrm{ns}$; it does not identify an independent replicate. SASA is expressed in $\mathrm{nm}^2$. The panels use different time and vertical scales.

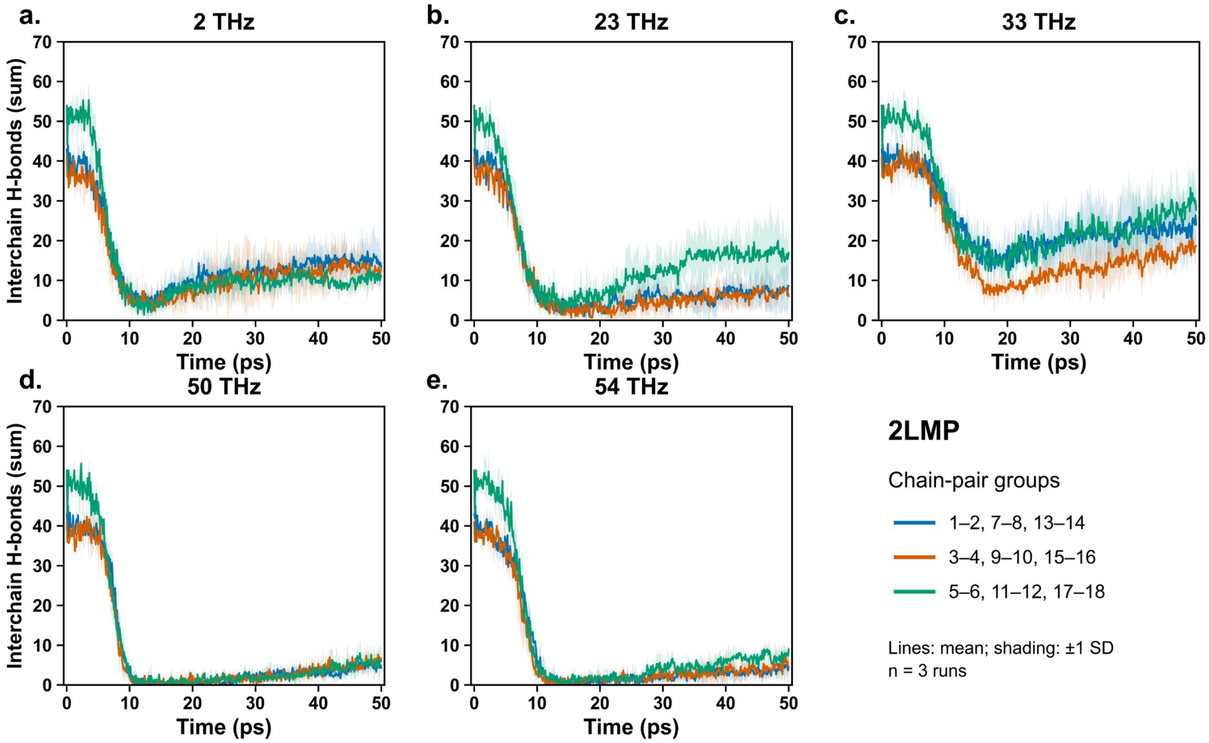


**Figure S4.** Interchain hydrogen bonds in three groups of 2LMP chain pairs under five pulse frequencies. Panels (a–e) correspond to 2, 23, 33, 50, and $54$ THz, respectively, at $E_0 = 2\ \mathrm{V\,nm^{-1}}$ with $x$ polarization. Blue, orange, and green denote the sums for pairs 1–2/7–8/13–14, 3–4/9–10/15–16, and 5–6/11–12/17–18, respectively. According to the plot legend, lines show the means and shading shows $\pm$ one standard deviation across three runs. The observation window is $0$–$50$ ps. The label $33$ THz denotes the $33.5$ THz input. Differences in remaining bond counts should be interpreted relative to the different initial counts of the groups.

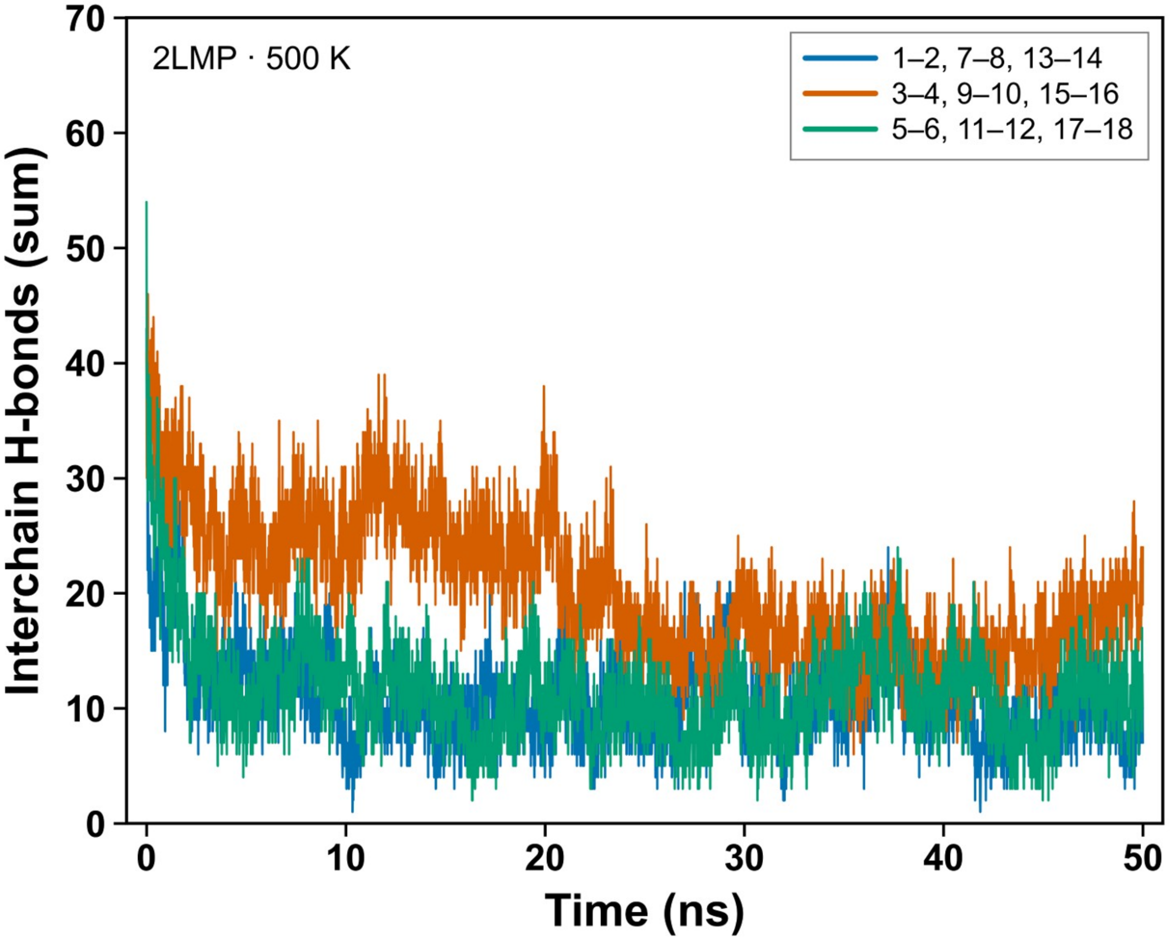


**Figure S5.** Interchain hydrogen bonds in the same three groups of 2LMP chain pairs during the 500 K field-free simulation. Blue, orange, and green identify the same pair groups as in Figure S4. The supplied panel displays 0–50 ns, distinct from the 0–50 ps pulse window and from the configured full heating duration in Table S2. This comparison characterizes interface reorganization under the heating protocol without matching pulse energy deposition or temperature history.

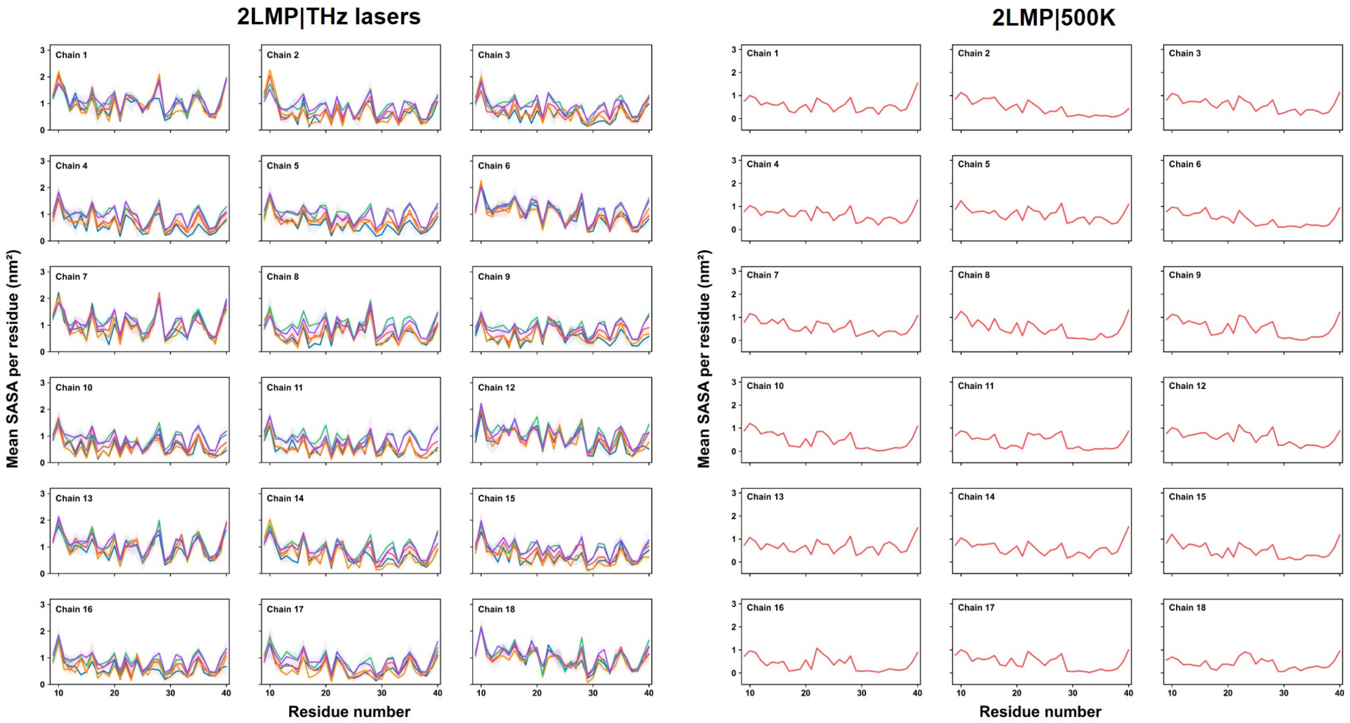


**Figure S6.** Residue-resolved mean SASA for all 18 chains of 2LMP. Left: time averages over $0$–$50$ $\mathrm{ps}$ for the 2, 23, 33, 50, and $54$ $\mathrm{THz}$ pulse conditions at $E_0 = 2\ \mathrm{V\,nm^{-1}}$ with $x$ polarization. The plot reports means and standard deviations across three runs, with the frequency colors defined in its legend. Right: the $500$ $\mathrm{K}$ panel, labeled as frame-weighted segment means over $0$–$244.22$ $\mathrm{ns}$ from one continuous run. Residue numbers are shown within each chain, and SASA is in $\mathrm{nm}^2$. The different averaging windows are retained explicitly.

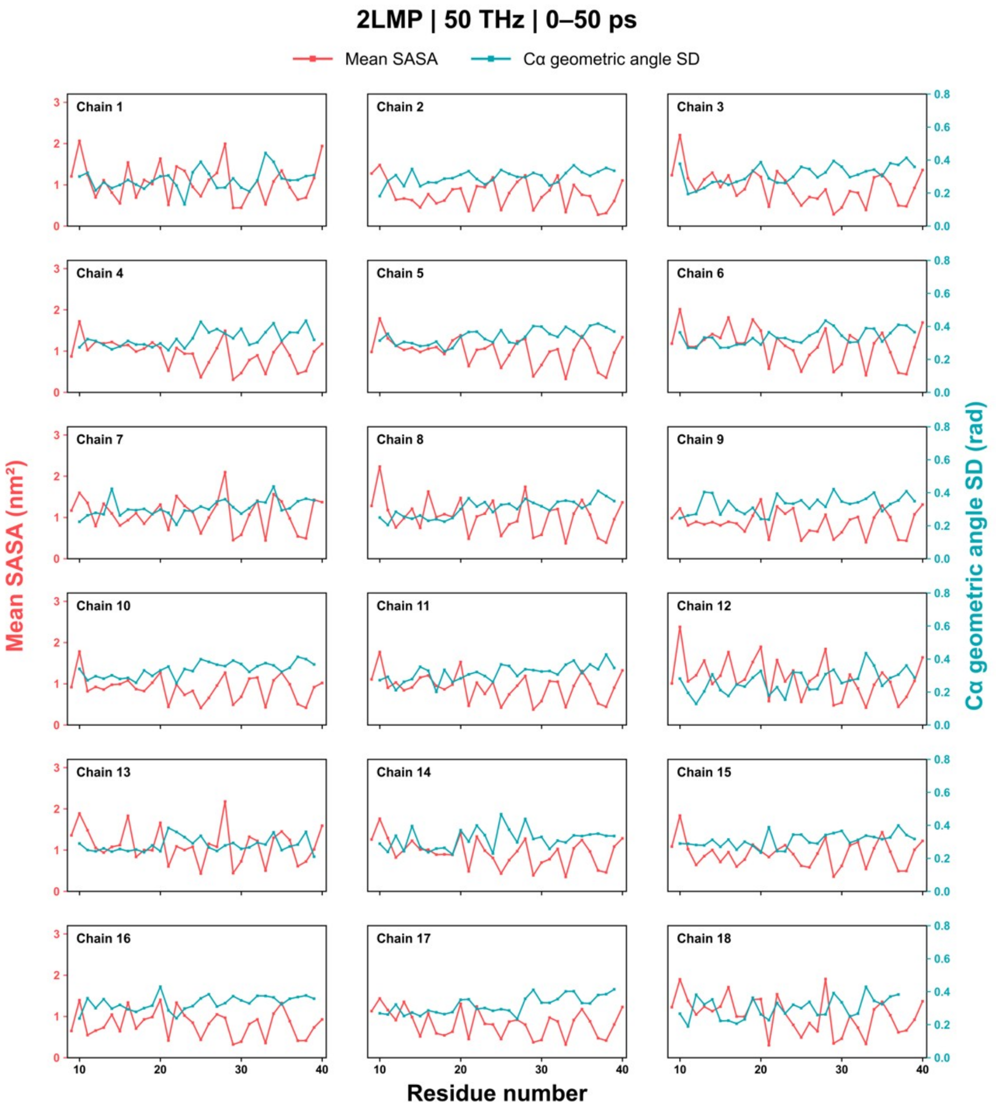


**Figure S7.** Residue-resolved solvent exposure and C$\alpha$ geometric-angle dispersion for the 18 chains of 2LMP under the $50\ \mathrm{THz}$ pulse at $E_0 = 2\ \mathrm{V\,nm^{-1}}$ with $x$ polarization. For each chain, red curves show mean SASA (left axis, $\mathrm{nm}^2$) and cyan curves show the temporal standard deviation of the geometric angle (right axis, rad) over $0$–$50\ \mathrm{ps}$. The comparison is exploratory. Whole-window angle dispersion includes conformational changes and does not quantify deposited vibrational energy.

Figure file to be supplied

figs8.jpg

**Figure S8.** Reserved: terahertz spectral comparison before and after laser excitation. Figure content, spectral definition, and caption will be added when this analysis is addressed.

Figure file to be supplied

figs9.jpg

**Figure S9.** Reserved: the two-threshold classification method associated with main-text Figure 6. The operational criteria, analysis windows, uncertainty estimation, and explanatory figure will be added when this analysis is addressed.

#### Supplementary references


[1] Anant K. Paravastu, Richard D. Leapman, Wai-Ming Yau, and Robert Tycko. Molecular structural basis for polymorphism in Alzheimer's $\beta$-amyloid fibrils. *Proceedings of the National Academy of Sciences*, 105(47):18349–18354, 2008. doi: 10.1073/pnas.0806270105. URL https://doi.org/10.1073/pnas.0806270105.

[2] Yiling Xiao, Buyong Ma, Dan McElheny, Sudhakar Parthasarathy, Fei Long, Minako Hoshi, Ruth Nussinov, and Yoshitaka Ishii. A$\beta$(1–42) fibril structure illuminates self-recognition and replication of amyloid in Alzheimer's disease. *Nature Structural & Molecular Biology*, 22(6):499–505, 2015. doi: 10.1038/nsmb.2991. URL https://doi.org/10.1038/nsmb.2991.

[3] Lothar Gremer, Daniel Schölzel, Carla Schenk, Elke Reinartz, Jörg Labahn, Raimond B. G. Ravelli, Markus Tusche, Carmen Lopez-Iglesias, Wolfgang Hoyer, Henrike Heise, Dieter Willbold, and Gunnar F Schröder. Fibril structure of amyloid-$\beta$(1–42) by cryo–electron microscopy. *Science*, 358(6359): 116–119, 2017. doi: 10.1126/science.aao2825. URL https://doi.org/10.1126/science.aao2825.

[4] William L. Jorgensen, Jayaraman Chandrasekhar, Jeffry D. Madura, Roger W. Impey, and Michael L. Klein. Comparison of simple potential functions for simulating liquid water. *The Journal of Chemical Physics*, 79(2):926–935, 1983. doi: 10.1063/1.445869. URL https://doi.org/10.1063/1.445869.

[5] Mark James Abraham, Teemu Murtola, Roland Schulz, Szilárd Páll, Jeremy C. Smith, Berk Hess, and Erik Lindahl. GROMACS: High performance molecular simulations through multi-level parallelism from laptops to supercomputers. *SoftwareX*, 1–2:19–25, 2015. doi: 10.1016/j.softx.2015.06.001. URL https://doi.org/10.1016/j.softx.2015.06.001.

[6] Kresten Lindorff-Larsen, Stefano Piana, Kim Palmo, Paul Maragakis, John L. Klepeis, Ron O. Dror, and David E Shaw. Improved side-chain torsion potentials for the Amber ff99SB protein force field. *Proteins: Structure, Function, and Bioinformatics*, 78(8):1950–1958, 2010. doi: 10.1002/prot.22711. URL https://doi.org/10.1002/prot.22711.

[7] Ulrich Essmann, Lalith Perera, Max L. Berkowitz, Tom Darden, Hsing Lee, and Lee G. Pedersen. A smooth particle mesh Ewald method. *The Journal of Chemical Physics*, 103(19):8577–8593, 1995. doi: 10.1063/1.470117. URL https://doi.org/10.1063/1.470117.

[8] H. J. C. Berendsen, J. P. M. Postma, W. F. van Gunsteren, A. DiNola, and J. R. Haak. Molecular dynamics with coupling to an external bath. *The Journal of Chemical Physics*, 81(8):3684–3690, 1984. doi: 10.1063/1.448118. URL https://doi.org/10.1063/1.448118.

[9] GROMACS Development Team. *GROMACS 2020.4 documentation: Molecular dynamics parameters*, 2020. URL https://manual.gromacs.org/documentation/2020.4/user-guide/mdp-options.html. Version 2020.4; accessed 4 October 2026.
